# Psychedelics relax predictive processing in the post-acute period by remodeling cortico-cortical feedback circuits

**DOI:** 10.1101/2024.07.03.601959

**Authors:** Chloe L. West, Fumiyasu Imai, Georgia Bastos, Molly A. Hornick, Lital Rachmany, Annabel Duran, Samen Nadeem, David Ricci, Anna M. Rader Groves, Joseph A. Wargo, Neil Van Leeuwen, Henry Sershen, K. Yaragudri Vinod, Darcy S. Peterka, Jordan P. Hamm

**Affiliations:** Neuroscience Institute, Georgia State University, Petit Science Center, 100 Piedmont Ave, Atlanta, GA 30303, USA; Department of Psychiatry, New York University Grossman School of Medicine, One Park Ave, New York, NY 10016; Emotional Brain Institute, Nathan S. Kline Institute for Psychiatric Research, 140 Old Orangeburg Rd, Orangeburg, NY 10692; Department of Philosophy, Florida State University, Dodd Hall, 151, 641 University Way, Tallahassee, FL 32304, USA; Schizophrenia Research, Nathan S. Kline Institute for Psychiatric Research, 140 Old Orangeburg Rd, Orangeburg, NY 10692; Child & Adolescent Psychiatry, New York University Grossman School of Medicine, One Park Ave, New York, NY 10016; Mortimer B. Zuckerman Mind Brain Behavior Institute, Columbia University, New York, NY 10027, USA; The Kavli Institute for Brain Science, Columbia University, New York, NY 10027, USA; Neuroscience Institute, New York University Grossman School of Medicine, One Park Ave, New York, NY 10016, USA

**Keywords:** psychedelic, mismatch negativity, oddball, saccades, anterior cingulate area, LSD, psilocybin, 5-MeO-DMT, predictive coding

## Abstract

Serotonergic psychedelics (e.g., psilocybin, LSD) have potential to treat psychiatric disorders, with therapeutic effects lasting days to weeks after a single dose. Prominent theories suggest that psychedelics have a lasting effect on hierarchical brain circuits, reducing top-down influence on information processing to facilitate an unbiased, bottom-up reassessment of the world, but direct and concrete evidence for such an effect is lacking. Here we directly tested this hypothesis in both humans and mice, assessing predictive processing in the fronto-visual system in the days after a single psychedelic exposure. Individuals who recently (<3 weeks) used 5-HT_2A_Receptor agonist psychedelics (psilocybin, LSD) were assessed via electroencephalography (EEG) and electrooculography recordings during a saccadic prediction task and compared to age- and sex-matched non-users. Compared to non-users, recent psychedelic users produced fewer fast saccades and less suppression of EEG delta/theta power to predictively presented stimuli, pointing to a disruption of predictive processing. These changes correlated with time since psychedelic use and were replicated in a second cohort taking a different serotonergic psychedelic (5-MeO-DMT). Direct recordings of primary visual cortex (V1) in mice administered psilocybin (1 mg/kg) evinced a similar loss of predictive suppression 24-hrs after the dose. This coincided with weakened top-down modulation of V1 from anterior cingulate area (ACa), a subregion of medial prefrontal cortex, along with clear spine growth in ACa neurons that project to V1. These results suggest that psychedelic-induced neural plasticity serves to reorganize feedback circuits in the cortex and relax top-down influence on bottom-up sensory processing – an effect that persists beyond the acute exposure period and may underlie a therapeutic window.

---

Serotonergic psychedelic compounds such as psilocybin, lysergic acid diethylamide (LSD), and 5-methoxy-N,N-dimethyltryptamine (5-MeO-DMT) have garnered interest in psychiatry for their potential to treat mood, anxiety, alcohol and substance use disorders (*1*–*5*). Elucidating the precise biological mechanisms underlying these therapeutic effects remains a central goal of basic and clinical research.

Like many antidepressant treatments (*6*), psychedelics stimulate plasticity in the mammalian medial prefrontal cortex (mPFC), changing neural connectivity via growth of dendritic spines (*7*–*9*). While the molecular mechanisms linking psychedelics to this spine growth are coming into focus (*10*, *11*), the functional consequences of such vast structural plasticity after psychedelic use are still unclear, particularly with regard to the circuit-level changes linking broad synaptic plasticity to lasting impacts on perception, cognition, and mood.

One reason for this gap is the lack of paradigms that both track the neuroplastic effects of psychedelics *and* can be translated to animal models for in depth mechanistic research. The head-twitch response in mice is widely used as a correlate of psychedelic effects, but, as an acute behavior that is observed only in rodents (*12*), it lacks validity as a clear marker of therapeutic and neuroplastic effects. Self-report and symptom scores in humans track the enduring effects of psychedelic medicine in clinical trials (*2*, *13*), but a) such measures are challenging to meaningfully translate to mice, b) may not cleanly map on to neural circuits, and c) suffer from the potential influence of bias due to inherent challenges with placebo control in such clinical trials (*14*).

Visual predictive processing is a sensori-cognitive function that translates well across species (*15*–*19*) and involves hierarchical brain circuits that are robustly engaged by serotonergic psychedelics (*20*). Conceptually, predictive processing refers to the influences of context and expectations on brain responses to sensory input by means of ongoing top-down modulation from higher brain areas to lower, often sensory, areas (*21*, *22*). Marrying this framework with psychedelic medicine, the Relaxed Beliefs Under pSychedelics (ReBUS) hypothesis posits that psychedelics exert their therapeutic effects (e.g., in depression) by altering predictive processing (*23*): by weakening the top-down influence of pathological biases, beliefs, and schema on information processing, allowing for a therapeutic reassessment of such biases in light of new experiences.

Although this theory refers mainly to higher order emotional and cognitive states - aspects which are difficult to translate to rodent research - it is plausible that psilocybin-induced changes apply to hierarchical brain circuitry more broadly, which include top-down cortico-cortical circuits that modulate sensory processing, a system that might be simpler to translate to mice (*16*, *24*, *25*). Further, mouse anterior cingulate area (ACa) is an mPFC subregion previously shown to exhibit spine growth after psilocybin (*7*, *11*), and ACa strongly modulates information processing in a predictive manner via feed-back projections to primary visual cortex (V1) (*15*, *26*).

Some past studies have suggested that basic predictive processing (mismatch negativity, [MMN] and P300 EEG potentials from visual and auditory oddball paradigms) is altered acutely during a psychedelic dose. Specifically, “deviance detection” – i.e., augmented brain responses to unexpected stimuli – is blunted during a trip (*27*–*30*). However, psychedelics also have dramatic effects on perception in this acute period, including large reductions in visual response gain in cortical neurons (*31*) and dramatic impacts on broader brain dynamics (*32*). Thus, studies of acute effects of psychedelics on brain function have limited utility for understanding the functional consequences of structural plasticity, which does not emerge until at least a day after a dose (*7*).

We studied individuals who had voluntarily taken a self-reported psychoactive dose of 5-HT_2A_ agonist psychedelics (psilocybin or LSD) in the prior 3 week period outside of a clinical setting. We recorded saccade latencies to predictable versus deviant target locations (saccadic prediction task; SPT), and electroencephalography (EEG) during the SPT and a passive visual oddball task. We found that recent psychedelic use correlated with reduced influence of predictability on reaction times and cortical responses to visual targets. Non-users produced accelerated eye-movements and reduced brain responses to stimuli in predictable locations (as expected), while recent psychedelics users did not. These effects correlated with time since acute drug exposure and were replicated in a second group of adults who took a different serotonergic psychedelic, 5-MeO-DMT.

Twenty-four hours after psilocybin administration, mice exhibited similarly altered visual predictive processing, showing reduced predictive suppression of neural responses to salient yet predictable stimuli in V1. Further, mice given psilocybin also exhibited weaker top-down modulation in V1 from mPFC (ACa).

We then examined and confirmed spine growth in the specific population of ACa intratelencephalic neurons (ITs) that project to V1. These results are consistent with the hypothesis that psychedelic-induced neural plasticity serves to reorganize in cortical feedback circuits, which impacts predictive modulation of sensory processing. This study outlines a potent translational paradigm for mapping circumscribed molecular, cellular, and systems-level mechanisms of psychedelics.

## Recent psychedelics users exhibit fewer short latency “express” saccades to predictably presented targets

Sixteen individuals who had recently taken psychoactive doses of 5-HT_2A_ receptor agonist psychedelics (PSY group; psilocybin [n=15] or LSD [n=1]) in the past 21 days (range 1 to 21 days) and sixteen age and sex matched comparison subjects (controls or CNT; see Table 1) were recruited for this study. PSY and CNT did not differ in major personality metrics (five-factor model (*33*)) or risk-taking behaviors, except for a slight increase in health/safety risk taking in the PSY group (t(30)=2.10, p=.04). Three subjects in the CNT group had used psychedelics previously at some point during their lifetimes, but were not group-level outliers in any of the major effects reported. The PSY group differed from the CNT group in the proportion of subjects using cannabis frequently (>4 times a month; Table 1), but the pattern of all major effects reported did not change when excluding the PSY participants with frequent cannabis use.

**Table 1:**
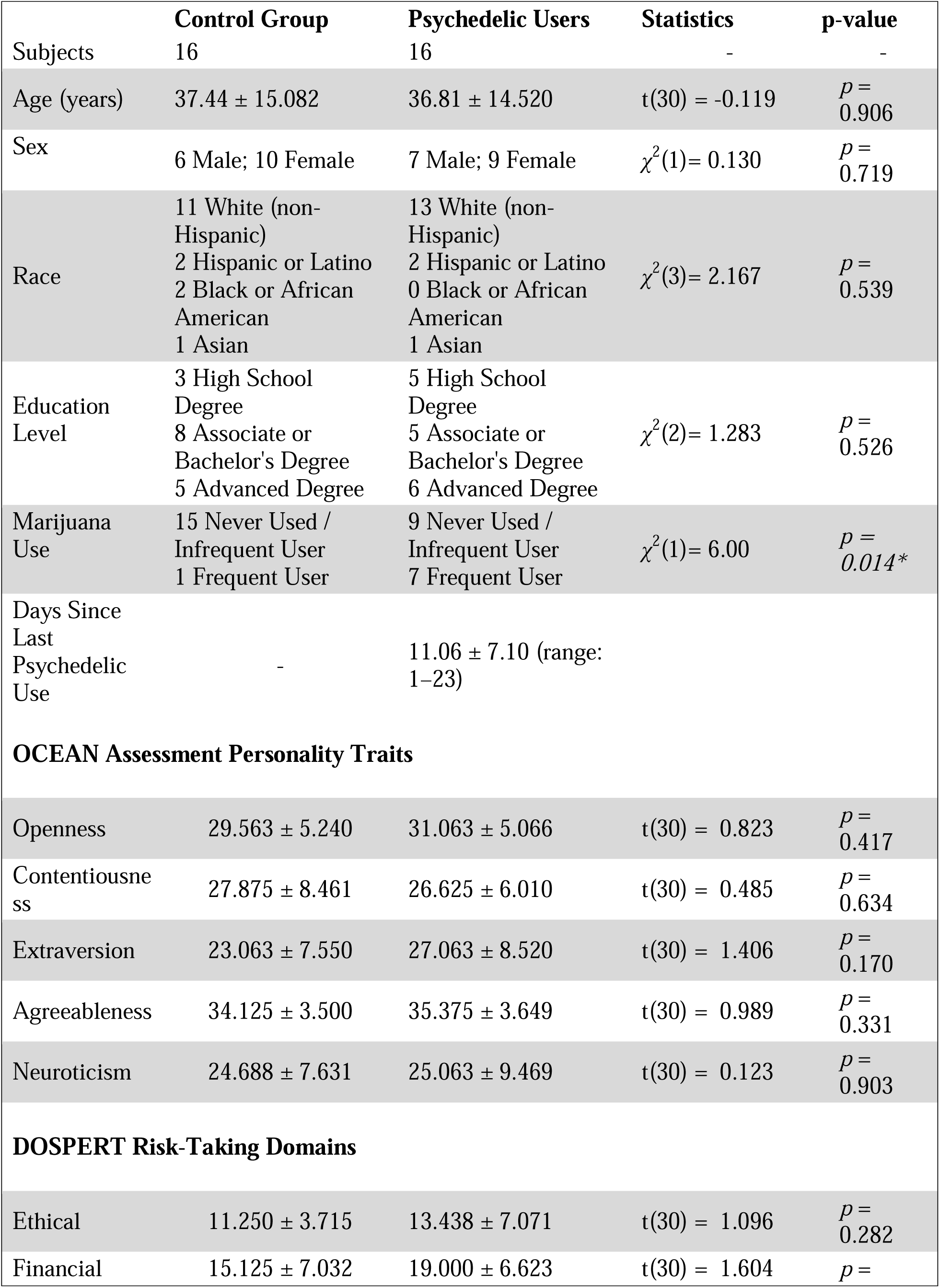
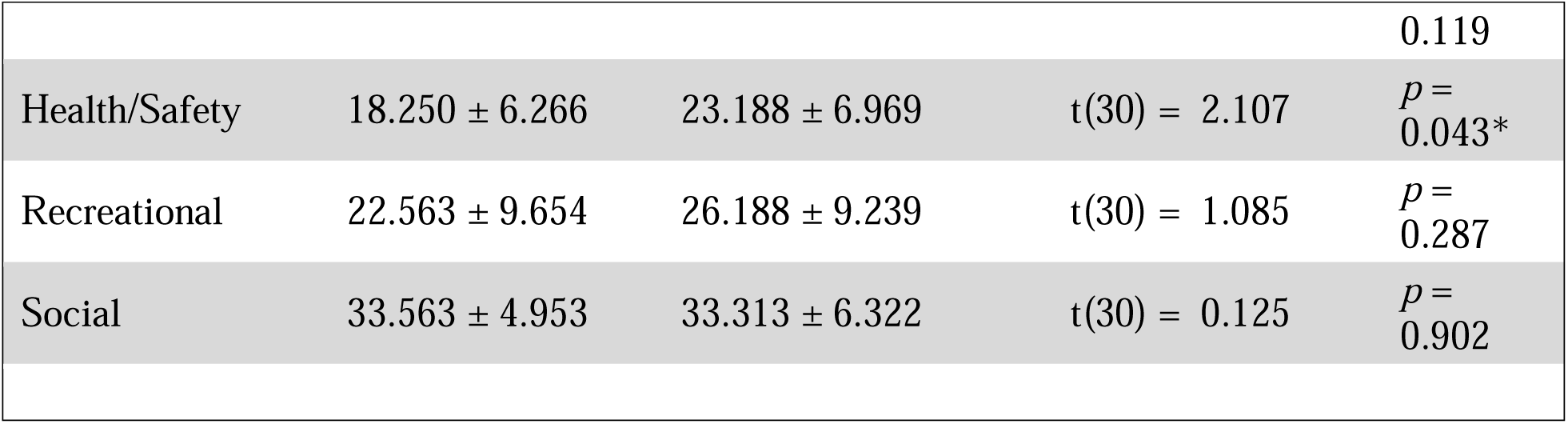
Demographic data, personality scores, and risk-taking among control and recent psychedelics-users. Group values represent means, +/- standard deviations.

First, we administered a SPT (*34*, *35*) to study predictive processing and updating after psychedelics use. The SPT requires subjects to fixate on a central cross (i.e., prestimulus) after which a star-shaped stimulus appears at one of eight locations (Fig. 1A). For the majority of trials (85%), the stimulus appears at a standard (predictable) location and is colored green or purple (“standard” trials). On a subset of trials (15%), the stimulus appears at a “deviant” (unexpected) location. On half of these deviant trials, the stimulus is orange and does not signal a change in the standard stimulus location (“deviant-no-update” trials). On the other half of the deviant trials, the stimulus is purple or green and signals a new “standard” stimulus location (“deviant-update” trials). This paradigm allows researchers to assess a) whether and how prior information about stimulus location affects behavior (i.e., response latencies to stimuli at expected or unexpected locations), and b) how individuals integrate new information to update behavior.

**Figure 1:**
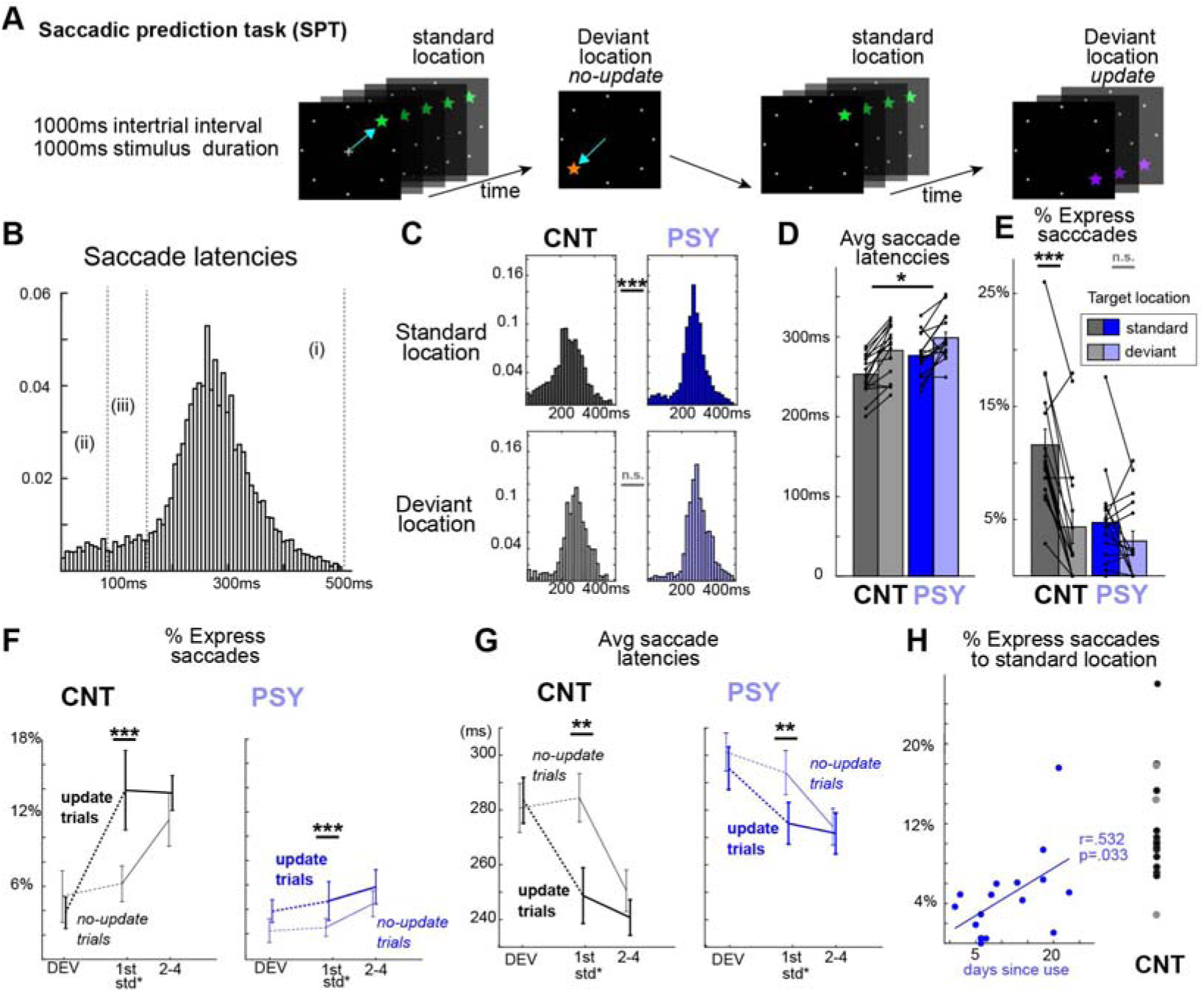
Recent psychedelic use reduces fast latency responses to predictable stimuli. A) The saccadic prediction task (SPT) involving saccadic targets in standard (predictable) and deviant locations, which were either informative of the upcoming targets (update) or not (no-update). B) Saccade latency histograms across all subjects and trial types, categorized as portions of (i) typical response latency saccades, (ii) putative anticipatory saccades, and (iii) fast- or express-saccades. C) Histograms by group and stimulus location. D) Average saccade latencies by group to standard and deviant target locations. E) Percentage of express saccadic responses (between 75 and 147ms) by group and target location. F) Percentage of express saccades and G) average saccade latencies for each group as a function of trial-type (x-axis) and deviant type (different lines). For F,G) *** reflects overall effect, across groups. H) Percentage of express saccades produced as a function of days since psychedelics use. Gray dots in the CNT group represent individuals with past use of psychedelics, but >11 months prior to the study. *p<.05, **p<.01, ***p<.001

We measured saccades through electrooculograms via two electrodes, placed just below and on the outer canthus of the left eye (Supplemental fig. S1). Saccadic reaction times for each participant were scored blind to group identity and trial type (see Methods). Overall, participants produced faster saccades to stimuli at standard (predictable) locations than to stimuli at deviant (or unexpected) locations (standard mean/std: 264.5ms/28.3ms; deviant: 290.3ms/31.3ms; F^location^(1,30)=44.8, p = 4 × 10^-7^). On average, CNT participants produced slightly faster saccades than PSY participants (CNT: 267.6ms/28.2ms; PSY: 287.2ms/24.4ms; F^group^(1,30)=4.4,p=0.04), but group differences in mean latency did not vary as a function of stimulus location (standard/expected location vs deviant location; F^interaction^(1,30)=0.97, p=0.33). However, comparison of the distributions of response latencies to different trial types suggested that predictability modulated response times differently between groups: CNT participants generated different types of saccades to predictable versus unexpected target locations (Fig. 1C; kstest=.069, p=0.008), while PSY did not (kstest=.038, p=0.34). CNT and PSY response distributions to the deviant stimulus locations were not different (kstest=.048, p=.33), while the CNT response distribution to the standard (expected) location differed dramatically from the PSY distribution (kstest=0.86, p = 2.5 × 10^-8^).

As illustrated in Fig. 1C, CNT participants produced an enriched proportion of short latency responses (<200ms) to targets in the standard location relative to the deviant location. This set of short latency saccades contains two subtypes: anticipatory saccades (ii in Fig. 1B) and express saccades (iii in Fig. 1B)(*36*). Saccades <75ms are considered anticipatory as this is the lower limit of visual afferent innervation of saccade-generating brainstem nuclei, and thus these saccades are non-visually driven (*37*). CNT and PSY did not differ in the overall percentage of anticipatory responses (Fig 1C; CNT: 6.3%/5.2%; PSY:4.1%/3.8%; F^group^(1,30)=2.2, p=0.15), nor was there a group by stimulus location interaction (F^interaction^(1,30)=0.16, p=0.69). On the other hand, express saccades are a subpopulation of very fast – *yet visually-driven* (non-anticipatory) – saccades with a mode around ≈110ms, separate from typical response latencies around 200ms (*36*, *38*, *39*). Such fast responses arise when stimulus location and timing is fixed or highly predictable (*40*). Express saccades reflect preparatory increases in excitability in visual and visuomotor regions driven by top-down modulation from frontal and parietal regions (*38*, *41*). The fact that express saccade proportion increases with practice (*40*) further suggests they encode learned spatiotemporal stimulus likelihoods.

We found group differences in the proportion of express saccades (Fig. 1C,E). Specifically, CNT participants produced a greater proportion of express saccades (75ms to 147ms; see methods for rationale) specifically to the standard stimulus (Fig. 1E). When compared with the PSY saccades, CNT saccades include a greater percentage of express saccades to stimuli in the standard location (CNT mean/std: 10.5%/4.3%; PSY: 4.3%/3.8%; t(30)=4.29, p=0.0002), but showed no difference from the PSY group in express saccades to the deviant location (CNT: 4.1%/5.5%; PSY: 3.0%/3.4.%; t(30)=0.687, p=0.49; F^interaction^(1,30)=13.53 p=0.0009). Each subject completed the SPT two times, and these same effects held for both runs (Supplemental fig. S2). Further, the overall percentage of express saccades to the predictable stimulus location was correlated with the time that had passed since the last use of psychedelics (Fig. 1H; r=0.532, p=0.033), suggesting that subjects begin to return to typical top-down control of saccadic circuitry after ≈7 days.

The SPT also allows for an assessment of how individuals use new information to update their behavioral strategies (*34*, *35*). The proportion of express saccades was mostly stable to stimuli in the standard location across trials except for stimuli immediately after the “no update” deviant (Fig. 1F). Subjects generated significantly fewer express saccades to the standard stimulus immediately after the “no update” deviant compared to the 2nd - 4th standards after the no-update (F^order^(1,30)=8.10, p=0.008). However, this was not true for the standard stimuli after the “update” deviant (F^order^(1,30)=3.02, p=0.093). This was also true for mean saccade latencies over all responses per subject (no-update: F^order^(1,30)=21.23, p=0.001; true update F^order^(1,30)=0.08, p=0.776). Interestingly, this pattern was present in both groups (Fig. 1G) and did not differ between groups (all interaction effects p>.198).

A slowing effect after the “no update” deviant is highly consistent with past studies using this paradigm and has been interpreted to reflect effects of updating (*35*). That is, a) participants in both groups made faster saccades overall to the standard stimulus occurring immediately after the updating deviant, suggesting rapid updating of internal models of probable target locations in both groups, but b) participants erroneously updated their expectations to “no update” deviants, effecting slower responses to the subsequent “standard” stimulus (which recovered quickly).

## Recent psychedelics users show less predictive suppression of early visual processing

Speeded responses to stimuli in predictable locations and times (*41*), and performance in the saccadic prediction task (SPT) in general (*34*), is thought to reflect top-down modulation of visuomotor processing based on context. If the lack of express saccades in PSY participants is driven by the attenuation of such top-down modulation, in line with the ReBUS theory (*23*), then the CNT group might also exhibit smaller cortical responses to stimuli in the standard location than to the deviant location while, the PSY group should show similar cortical responses to the stimulus in the standard location as to the deviant location. We recorded 32 channel electroencephalography (EEG-Biosemi) while subjects completed the SPT. Based on averaged event-related responses (ERPs) to stimuli in the time-voltage domain, PSY participants appeared to demonstrate greater cortical responses to all stimuli (Fig. 2). We analyzed the EEG activity in the time-frequency domain to increase signal to noise ratio (as the number of trials was ≈20 for some conditions), facilitate translation to animal models (*42*, *43*), and disentangle increases in signal power from phase-alignment across trials (*38*, *44*).

**Figure 2:**
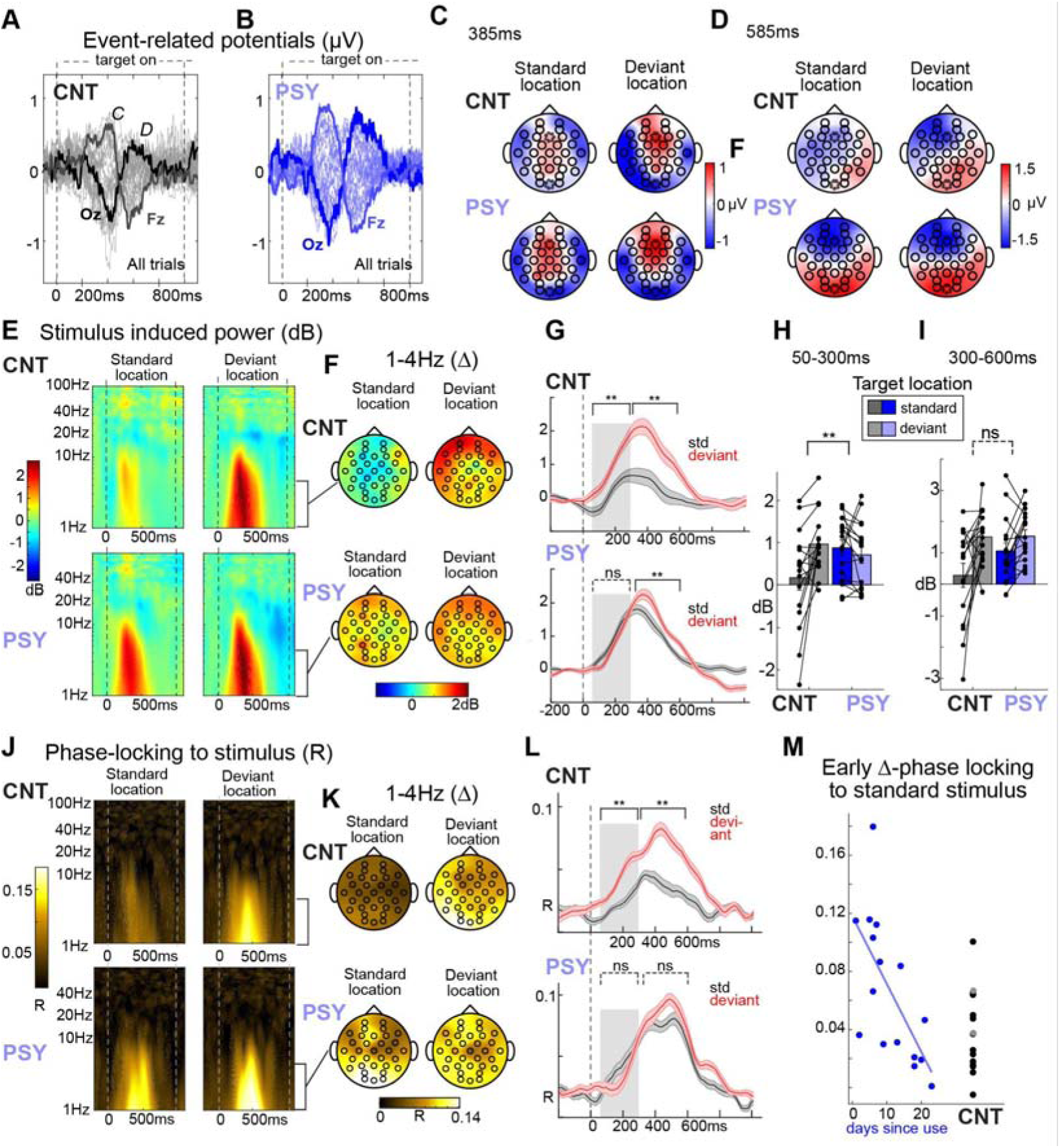
Psychedelics alter predictive modulation of visually evoked responses. A) Butterfly plot of event-related potentials to all stimuli (each line is one of 32 electrodes) averaged across all CNT and B) PSY subjects. C) Averaged scalp topography from first and D) second peaks in A,B. Fz and Oz indicated as dotted lines. E) Stimulus induced power, averaged over all electrodes, to targets in the standard location and the deviant locations for CNT (top) and PSY (bottom) subjects. F) Scalp topographies showing averaged delta-band power from 50 to 600ms post-stimulus onset. G) Time-course of averaged delta-band power from all electrodes. H) Bar plots showing early time window (50-300ms) and I) later time window (300-600ms) delta power from (I). J-L) same as E-G, but for inter-trial phase locking. M) Correlation of delta-band phase locking across all electrodes during the early period (50-300ms) with days since psychedelics use. **-p<.01

Consistent with our hypothesis, trial-averaged power (i.e., average single-trial power) to the onset of saccadic targets appeared weaker to stimuli in the predictable (standard) location as compared to stimuli appearing in deviant locations in CNT (Fig. 2E; averaged over all electrodes, 50-600ms post-stim onset), and this was generally true across the scalp in low frequencies (Fig 2F). We split analyses into early (50-300ms) and late (300-600ms) based on the rise and fall of the power waveform (Fig. 2G) as well as the established latencies of typically analysed early (e.g., N100, MMN) vs late (P300) evoked cortical activities (ERPs), respectively. We focused on traditional frequency bands (delta 1-4Hz, theta, 5-8Hz; alpha 9-13Hz; gamma 35 to 100 Hz), and separately examined effects of prediction (standard vs all deviants) and updating (deviant update vs deviant non-update).

For the early period, CNT showed strong predictive modulation, with much stronger delta band responses to deviant stimuli relative to predictable stimuli (Fig. 2G, H) while PSY showed no difference (F^interaction^(1,30)=9.25, p=.0049;, CNT-t(15)=3.052, p=.0081, PSY-t(15)=0.931, p=.3662). The same group by stimulus location interaction effect was present for early theta power (F^interaction^(1,30)=7.7, p=.0094; Supplemental fig. S3). For later processing periods, these group by stimulus location interactions were absent (delta– F(1,30)=2.29, p=.141; theta–F(1,30)=1.18, p=.285), as both PSY and CNT groups demonstrated augmented delta (Fig. 2I) and theta power (Supplemental fig. S3) to deviant stimuli. This suggests that later brain responses to contextually deviant stimuli – putative prediction-error signals or downstream “deviance detection” – were unaffected by recent psychedelics use.

Consistent with this interpretation, the degree of EEG phase-locking to stimulus onset was similarly impacted by psychedelic use. While single-trial power may reflect both stimulus-entrained and endogenously-driven neural processing of a stimulus, inter-trial phase-locking to the onset of sensory stimuli specifically captures stimulus-entrained signals and may better index bottom-up processing of sensory inputs (*44*, *45*). Frequency bands above delta did not show strong phase-locking factor (PLF) for either group, so our analyses focused on delta. CNT participants showed increased PLF to stimuli in the deviant locations compared to stimuli in the standard location in delta frequencies, while PSY did not exhibit this effect in early time ranges (50-300ms; delta: F^interaction^(1,30)=8.61, p=0.006, CNT-t(15)=5.296,p=.00009, PSY-t(15)=-.805, p=.4335; Fig. 2J-L). Similar to power, groups showed more modest differences in late delta PLF prediction effects, with only trend-level significance (F^interaction^(1,30)=3.93, p=0.06). Finally, the magnitude of early PLF to predictable stimuli was inversely correlated with the number of days since psychedelics had last been taken (r=-.670, p=0.0045; Fig. 2M), in line with the effect on percent of express saccades to this stimulus.

We did not observe any significant effects of update type (non-updating vs updating deviant) for either group for any frequency band for early or late time periods, for PLF or power (Supplemental fig. S3), although there was a trend toward higher gamma power in the late period for updating deviants, present in both groups F^stim-location^(1,30)=2.64, p = .11). Other frequency bands (alpha, beta, gamma) showed no main or interaction effects involving group or stimulus for either timebin for PLF or power.

In sum, these EEG results concord with the behavioral results and suggest that only specific aspects of visual predictive processing are altered after psychedelics use. Late processing of prediction errors (Fig. 2G,I) and context updating are not altered after psychedelics (saccade behavior –Fig. 1F-G). On the other hand, top-down modulation of early sensory processing – *i.e., the degree to which learned patterns modulate early or preparatory processing of subsequent stimuli* – is weakened for days after a dose.

## Alterations in visual predictive processing generalize to other serotonergic psychedelics

We additionally recruited 15 subjects who had recently (<24 days) taken a different serotonergic psychedelic compound, 5-MeO-DMT. Like psilocybin and LSD, 5-MeO-DMT involves an intense psychedelic experience involving altered perception, albeit with strong affinity for 5-HT_1A_ in addition to 5-HT_2A_ receptors (*4*), and has demonstrated efficacy in clinical trials for depression (*46*). Our 5-MeO-DMT group did not significantly differ from CNT in any demographic, lifestyle, or risk-taking behaviors, and only showed a moderately increased openness compared to the CNT group (D=0.81; Supplemental Table S1).

We focused on the major effects that differentiated the PSY group from CNT in figures 1 and 2. Effects are presented in Supplemental fig. S4. Like the PSY group, the 5-MeO-DMT group showed a reduction of express saccades as a function of location compared to CNT (fewer to the predictable location; F^interaction^(1,29)=8.32, p=.007; fig. S4C). Further, like in the PSY group, there was a group by stimulus location interaction on early delta power (F^interaction^(1,29)=7.75, p=.0094; fig. S4H) and phase locking (F^interaction^(1,29)=15.32, p=.0009; fig. S4M) in the 5-MeO-DMT group, whereby the 5-MeO-DMT group did not show reduced early responses to the stimulus in predictable locations. Later power effects of stimulus location did not differ from CNT (F^interaction^(1,29)=1.83, p=.186; fig. S4I), as was seen in the PSY group, but later PLF did (F^interaction^(1,29)=9.65, p=.0042; fig. S4N), again similar to the PSY group, potentially reflecting the differential sensitivities of PLF vs power to stimulus-locked neural activities. Like the PSY group, the 5-MeO-DMT group showed evidence of intact updating in their behavioral metrics, as they did not differ from CNT in a slowed recovery from a non-updating deviant (F^stims-after-deviant^(1,29)=7.76, p=.0093; F^stim-by-group-Interaction^(1,29)=0.99, p=.327) or in a fast recovery after an updating deviant (F^stims-after-deviant^(1,29)=0.48, p=.492; F^stim-by-group-Interaction^(1,29)=0.35, p=.558). Effects in both 5-MeO-DMT and PSY were present regardless of sex or age (Supplemental figure S5).

In sum, a second group of subjects taking a different serotonergic psychedelic exhibited the same effects on behavioral and neural indices of predictive modulation of early visual processing, along with unaffected indices of late prediction error processing or context updating.

## Psilocybin alters predictive processing in primary visual cortex in mice

To directly test whether a dose of serotonergic psychedelics can alter visual predictive processing in the post-acute period, and to determine whether these effects are present in sensory processing regions, as suggested by the early latency effects in figure 2, we administered to mice a dose of psilocybin (PSL (from Usona Institute); 1mg/kg; I.P.; n=12: 8 male, 4 female) or saline (SAL; n=7; 4 male, 3 female). This dose was sufficient to induce a canonical head-twitch response in a subset of mice (later used for histology), confirming psychedelic-like effects (Fig. 3A,B). Local field potentials were recorded in layer 2/3 of primary visual cortex (V1; Fig. 3C) as previously reported (*16*).

**Figure 3.**
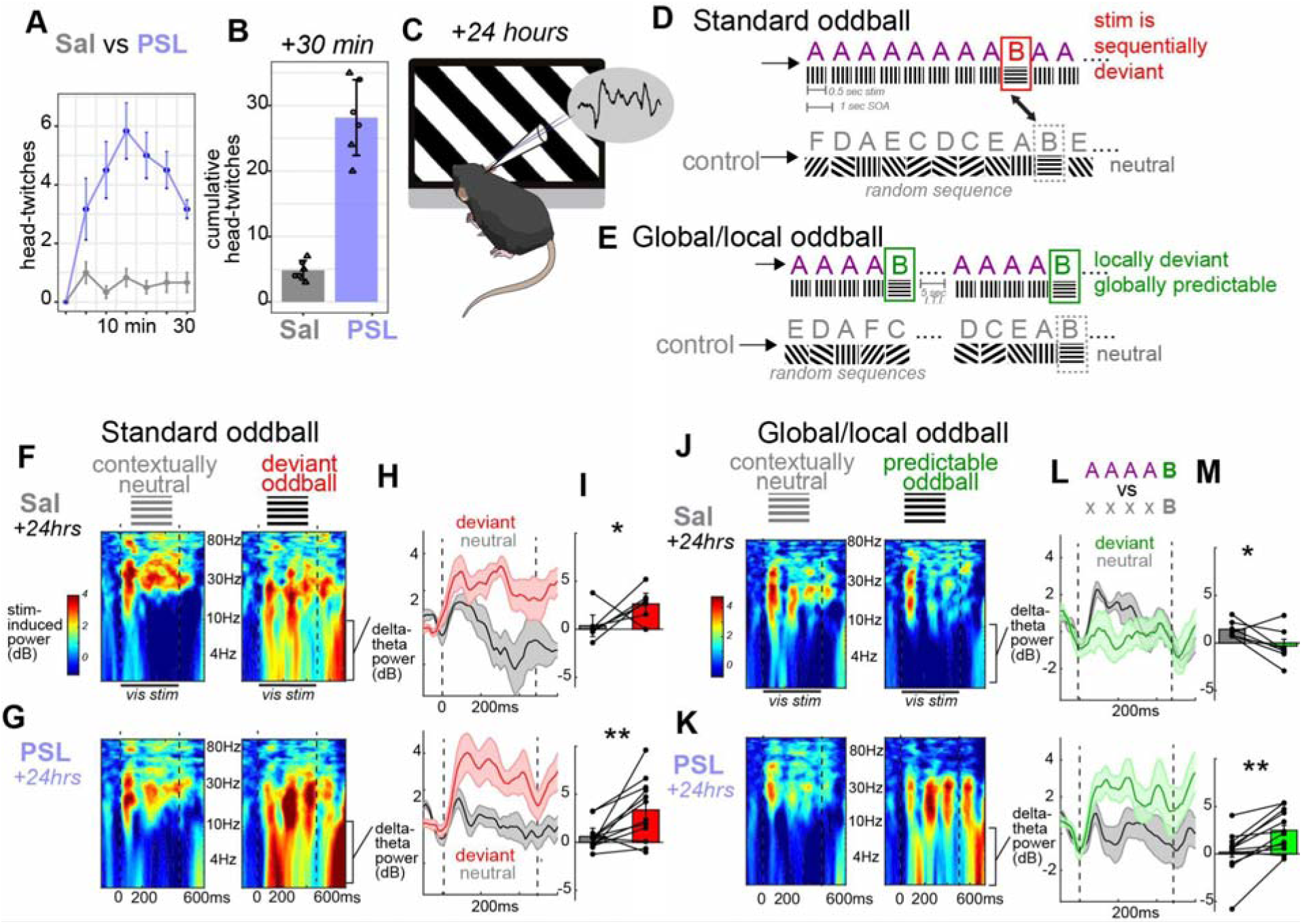
Predictive suppression in primary visual cortex is diminished 24 hours after a dose of psilocybin. A,B) Canonical head-twitch responses were present in the first 30 minutes after a dose of 1mg/kg psilocybin (PSL), relative to saline treated mice (Sal). C) 24-hours after a dose, local field potentials were measured in awake mouse primary visual cortex (V1) during visual stimulation using full-field visual grating stimuli. D) A standard oddball paradigm was presented (with deviants occurring at random times) and compared to a control sequence where stimulus orientation was random. E) A global-local oddball paradigm was presented, where oddball orientations were always presented after 4 redundants (AAAAB; 35 repeats), and was compared to a control sequence consisting of 5 randomly selected orientations. In (D) the B orientation occurs at unexpected times, while in (E), the B orientation occurs at a predictable time, though in both cases, it occurs once every 8-10 seconds. F) Stimulus induced power spectra from saline treated and G) psilocybin treated mice from the standard oddball in (D). H,I) Average power from 2-10Hz (delta/theta) from 50-300ms (early time window) shows suppression of deviance detection in both Sal and PSL mice. J-M) Same as F-I, but during the global/local paradigm in (E), when the B stimulus is temporally predictable. *p<.05; **p<.01.

Twenty four hours after the dose (post-acute period), we examined visual predictive processing in awake head-fixed mice using a “standard” oddball paradigm (*42*, *47*) and a “global-local” oddball paradigm (*15*, *48*). Stimuli were full-field 100% contrast drifting squarewave gratings (0.08 c.p.d.; moving at 2 Hz; oriented at 8 possible angles, 0 to 157.5 deg; 500ms stimuli, 500ms I.S.I). In the standard oddball, one stimulus orientation (i.e., 90-deg bars or “A” stimuli in Fig. 3D) is presented repeatedly, and is occasionally interrupted at random times by a different “deviant” orientation (i.e., horizontal bars (0-deg) or “B” stimuli Fig.3D). Responses to the B orientation in this mostly predictable context are then compared to responses to B when it occurs during a more neutral context in which B is equally likely to appear as any other orientation – a “many-standards control” sequence (Fig.3D) to isolate “deviance detection” (i.e., augmented cortical responses to unexpected stimuli – a form of visual prediction error (*15*, *49*, *50*). In contrast, during the “global-local” paradigm (Fig. 3E), B stimuli come after a sequence of A stimuli as well, yet always appear at the end of a 5 stimulus sequence (AAAAB; Fig. 3E). Thus, in the global-local paradigm, B is locally deviant but globally predictable (i.e., sequentially expected), while in the standard oddball paradigm, B occurs at unpredictable times. In both cases, and in the control runs, B orientations occur at approximately once every ≈8-10 seconds. This approach provided us with the advantage of needing limited training (mice exhibited deviance detection and suppression on the first day of exposure, 1 hr prior to dosing; Supplemental fig. S6), and the ability to compare an equally salient stimulus (B) in both conditions (similar to a saccadic target in the human run) that is temporally predictable in one case (the global-local oddball) but not the other (the standard oddball).

Global predictive processing implies that stimuli are processed in the context of learned probabilities or broader spatiotemporal patterns beyond local features (e.g., immediately previous or concurrent stimulus differences), requiring top-down modulation of sensory processing from higher brain regions (*21*, *22*, *51*). So, deviance detection to the B during the standard oddball paradigm, but not to the B during the global-local paradigm, would suggest that global predictive processing is intact. Deviance detection to both would suggest that local dynamics are intact, but global predictive processing is missing. Given that the latter requires top-down modulation from higher brain regions, we hypothesized that, like our human findings and in line with the ReBUS hypothesis (*23*), predictive processing in V1 would be altered after psychedelics.

At this baseline, groups did not differ, with both exhibiting deviance detection only during the standard oddball paradigm (Supplemental fig. S6). Then, 24 hours after treatment (Sal or PSL), we observed a paradigm (standard vs global-local) by context (deviant vs control) by treatment interaction (F(1,67)=5.35, p=.024) on delta-theta power (2-10Hz) during the same early response period above (50-300ms). The Sal group exhibited a paradigm by context interaction (F(1,23)=14.489, p=.001) driven by deviance detection during the standard oddball (t(6)=2.03,p=.044) and deviance *suppression* during the global-local paradigm (t(6)=-2.449, p=.049), suggesting that responses to a locally deviant –but otherwise salient –stimuli were bidirectionally modulated by context (Fig. 3F,H,I,J,L,M). Conversely, the PSL group showed strong deviance detection to both deviants, regardless of context (standard t(11)=3.14,p=.009; global-local t(11)=4.397, p=.001) and no paradigm by context interaction (F(1,43)=0.216, p=.645; Fig. 3G-I,K-M).

During the global-local paradigm, we also included catch trials with a deviant 5th item in the sequence other than B, as previously described (*15*). On 6 of the 70 sequence repeats (trials) the sequence was AAAA-C (a global and local deviant, where A was 90 degrees and C was 45 degrees) and on another 6 trials, the sequence was AAAA-A (a local redundant but a global deviant; all 90 degrees). These results are presented in supplemental figure S7. The SAL group exhibited significant deviance detection to both the C deviant (t(6)=3.798, p=.0045) and the A deviant (t(6)=3.097, p=.011), suggesting strong global predictive processing in the SAL group. On the other hand, the PSY group only displayed significant deviance detection to the C deviant (t(11)=3.324, p=.0034), but not to the A deviant (t(11)=1.251, p=.118).

Therefore, psilocybin had a lasting post-acute impact on context-based suppression of salient but predictable stimuli in the visual system, aligning with effects seen in human participants during a predictive saccade task (Fig. 2). In a subset of human participants, we also collected the standard oddball paradigm, and results confirm that basic deviance detection was intact after psychedelics, despite a loss of predictive suppression during the SPT task (Supplemental fig. S8). The fact that deviance detection effects were present in V1 a day after psychedelics regardless of global context suggests that local predictive processing might dominate in visual cortex in the period after psychedelics, due to weakened top-down modulation from long-range projecting neurons in higher brain regions.

## Psilocybin alters top-down modulatory circuits from mPFC to V1

Predictive processing in V1 depends on top-down feedback from higher brain areas, including ACa (*26*, *49*). To more directly test whether changes in visual predictive processing observed after psychedelics (Figs. 1-3) reflect alterations in the balance of top-down modulation versus bottom-up information processing in the cortex (*23*), we also recorded local field potentials from ACa along with V1 and calculated non-parametric Granger causality analysis across frequencies (1-40Hz) to assess directed functional connectivity between these nodes. We analysed whether top-down (ACa-to-V1) and/or bottom-up (V1-to-ACa) connectivity differed after psilocybin during the global-local paradigm, where predictive suppression was observed. We compared top-down vs bottom-up Granger coefficients at 11-12ms lags (based on past work and estimated conduction delays (*16*)) in 3 bins – first stimulus, middle stimulus, final stimulus and focused on the delta-band, where the top-down influence was maximal (Figs. 4B, S6I). We found that, 24 hours after psilocybin, there was an alteration in the balance of top-down vs bottom-up connectivity in this circuit in the delta band (F^treatmentXdirection^(1,95)=8.527, p=.004), regardless of time bin, suggesting broad weakening of top-down modulation after psychedelics (Fig. 4B,C). Groups did not differ at baseline (Supplemental fig. S6; F(1,114)=0.710; p=.401).

**Figure 4.**
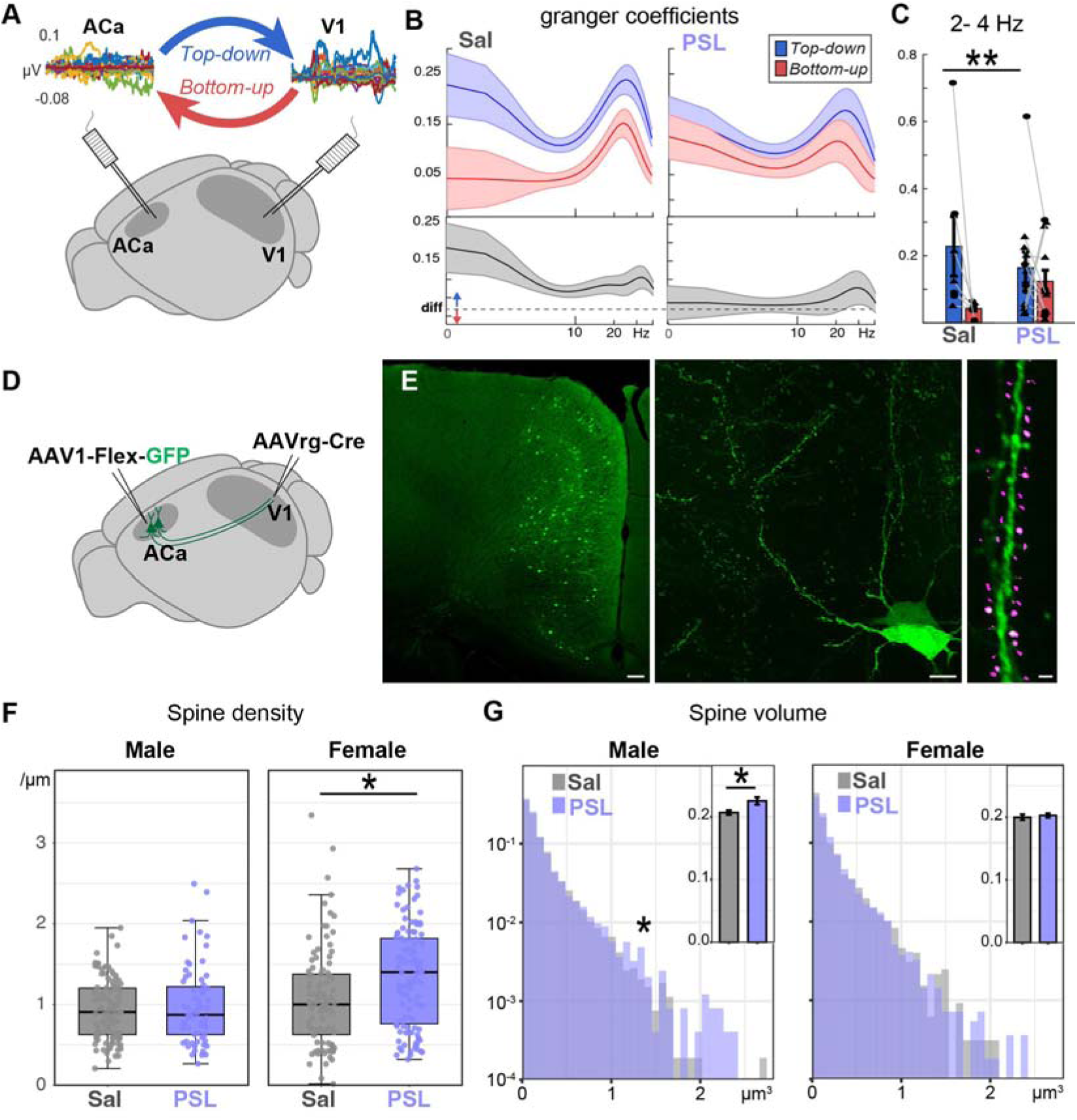
Psychedelics alter feedback circuit function and structure in the cortex. A) Local field potential recordings in ACa and V1 in awake mice during visual stimulation 24 hours after dose. B) Top-average spectra of top-down and bottom-up Granger coefficients (directed influence from ACa to V1 or V1 to ACa, respectively) for saline control (left) vs psychedelic groups (right). Bottom - top-down minus bottom-up spectra. C) Average GC values for all stimulus/ITI time-bins. Each dot is one mouse, triangles are males and circles are females. D) Retrograde labelling of ACa neurons that project to V1 with GFP and E) Images of labelled neurons at different scales. Right panel demonstrates scoring with RESPAN (Magenta). Scale bars: 100µm (left), 10µm (middle), 1µm (right). F) Spines per micrometer of dendrite. Each dot reflects a dendritic segment. G) Average spine volume histograms with inset bar plots displaying overall means (across spines) and 95% confidence intervals. *p<.01.

Recent work in mice has shown that psilocybin induces plasticity in ACa pyramidal neurons (*11*), including both pyramidal tract (PT) and intratelencephalic (IT) projection neurons. While the plasticity in the former type, PTs, which project to subcortical areas, has been linked to shifts in stress-related behaviors (*11*), the consequences of spine growth in IT-type neurons are unknown. Crucially, the ACa IT population could include feedback neurons that project back to V1 to support predictive processing (*26*). We sought next to determine whether this specific subpopulation of ACa neurons that project to V1 exhibit spine growth after psychedelics. Using a retrograde-Cre labelling approach (Fig. 4D), we expressed GFP in ACa neurons that project to V1, and then fixed tissue 24-hrs after the saline or psilocybin dose. We utilized the RESPAN Toolbox (*52*) to quantify spine density in n=384 dendritic segments and spine volume in n=16,597 spines from 6 mice treated with psilocybin (3 male/3 females) and 6 mice treated with saline (3 male/3 females). We found that psilocybin generally induced spine growth in both sexes of mice, however, there were sex by drug effects in both metrics, consistent with some past reports in mPFC after psilocybin (*7*). Spine density increased only in females (Fig.4F; F^sexXgroup^(1,378)=5.006, p=.0258) while spine volume increased only in males (Fig. 4G; F^sexXgroup^(1,16591)=18.242, p=.00002). Although some past reports have identified sex-dependent dose-response effects of psilocybin on alcohol consumption (*5*), we did not observe significant sex by group interaction effects in any functional or behavioral effects humans or mice (figure S5), suggesting that these sex differences in structural metrics might reflect similar plasticity with equivalent downstream consequences in cortical circuits.

## DISCUSSION

A principal function of the neocortex is to process incoming sensory data in the context of concurrent and previous stimuli, perceived spatiotemporal patterns, motor outputs, and behavioral goals. Typically, a stimulus with contextually predictable features evokes smaller responses in sensory cortex than one with contextually deviant features, and subsequent reactions to that stimulus are facilitated by this predictability. Our study provides evidence for this in the comparison group of humans (Figs. 1, 2) and in mice treated with saline (Fig. 3). We then show that a dose of serotonergic psychedelics alters these predictive processing functions in the post-acute period, after the acute effects on perception have subsided (>24 hours). Specifically, in humans, responses to predictable yet salient stimuli (a saccadic target occurring in a repeated location) were no longer suppressed relative to contextually unexpected stimuli (a target in a novel location). In mice, a 1mg/kg dose of psilocybin also led, 24-hours later, to a similar loss of predictive suppression within primary visual cortex (V1). That is, when saline-treated mice were habituated to expect a change in visual stimulus orientation occurring at a specific interval, their visual cortices no longer responded strongly to this change. Yet, after psilocybin, this suppressive effect was lost. These effects coincided with weakened feedback modulation of V1 from neuronal populations in a higher brain area (ACa) — a region previously shown to support predictive processing through long range connections to sensory regions (*15*, *16*, *26*, *53*). These long range projection neurons in ACa also exhibited growth in dendritic spine number (females) or size (males; Fig. 4) after psilocybin.

These results concord with the hypothesis that psychedelics alter predictive processing in hierarchically organized brain networks (*23*), yet also emphasize these effects in the post-acute period, in the days after a dose (*54*). In a predictive processing framework (*21*, *51*), higher-order brain areas modulate sensory processing in lower areas in line with expectations, observed contextual regularities, and/or perceptual priors: suppressing responses to sensory inputs that fit with current perceptual “priors” (gleaned from past experience) and augmenting responses when they do not (prediction errors)(*22*, *51*, *55*). Consistent with this framework, oddball paradigms used here (*15*, *16*, *26*) (along with sensorimotor (*56*) and visuospatial paradigms (*49*) all highlight a central role of feedback inputs to V1 in predictive suppression and prediction errors (*57*). In the current study, the former – predictive suppression – was specifically disrupted after psychedelics, which coincided with decreased directional functional coupling (Granger causality) in the top-down direction of the ACa-V1 circuit. However, ACa IT neurons that mediate this connection showed an increase in spines in the current study, despite their decreased long-range influence. Consistently, past work on the mPFC to amygdala network (*58*) and other cortico-subcortical networks (*59*) have observed a similar post-acute weakening in connectivity in fMRI BOLD, despite the fact that synapses in mPFC PT neurons that project subcortically *increase* (*7*, *8*, *11*), like IT neurons in the current study.

We propose that the growth in spines in higher brain areas after psychedelics is the structural substrate of decreased precision in high-level priors, which could affect both subcortical and cortical processing. Specifically, it is possible that, as psychedelics induce a broad and non-specific increase in synaptic connectivity within mPFC, pre-existing mPFC ensemble patterns are partially lost in favor of a shift toward all-all connectivity. This serves to weaken effective connectivity on distal target regions (e.g. V1), which otherwise precisely integrate hardwired top-down inputs through local inhibitory (SST+ interneurons) and disinhibitory (VIP+ interneurons) microcircuitry (*16*, *24*). Altered engagement of these post-synpatic inhibitory circuits (by psychedelic reorganized top-down projections) may thereby weaken stimulus-specific inhibitory control of V1 neural populations. To test this model, future work should aim to directly block spine growth in the subset of neurons that project from ACa to V1 and determine whether this a) alters ACa-to-V1 effective connectivity (Granger causality), b) ACa ensemble membership, e.g., (*60*, *61*), and c) predictive processing in V1.

This study provides a candidate functional consequence of IT neuron plasticity after psychedelics. Past work in mice demonstrates that psilocybin increases spines in two major types of excitatory neurons in the mPFC – pyramidal tract neurons (PT) and intratelencephalic neurons (IT) (*6*, *11*). While the former projects mainly to subcortical areas, such as amygdala, thalamus, and brainstem, the latter projects mainly within the cerebral cortex, including lower sensory areas (*62*). In humans, amygdala responses to emotional stimuli are known to change in the day after a psychedelic dose (*63*) and mouse work has linked changes in depressive- and anxiety-like behaviors in mice to PT neuron plasticity in mPFC (*11*), providing a neural substrate. However, the impact of spine growth in IT neurons – which by many measures is equal or greater than PT neuron spine growth in mPFC – is not known. Here we confirm IT neurons within mPFC that project to V1 exhibit spine growth and suggest a functional impact of this plasticity: softened top-down modulation which weakens predictive suppression.

These results highlight the lasting impacts of psychedelics on top-down modulation in the brain – an effect that more broadly could help explain the therapeutic benefits of these compounds (*23*). Interestingly, psychedelics consistently increase cognitive flexibility in both healthy individuals (*32*) and individuals with major depression (*64*) in the post-acute period. Temporarily weakened top-down influence on information processing offers a potential mechanism underlying this cognitive flexibility. This mechanism, in theory, would allow new information to escape the influence of overly negative schema or beliefs (as is proposed in major depressive disorder (*65*) and update them. To confirm this possibility, future studies should aim to correlate visual predictive processing measures (such as those included in this study) with treatment response in clinical trials among various psychiatric populations. If confirmed, it may help recenter future research on enhancing IT neuron plasticity. On the other hand, it is possible that the core therapeutic effects of psychedelics rests in their modulation of PT neurons (*6*, *11*), while IT neuron plasticity may have differential, mostly perceptual, impacts, and thus, may be dispensable for the lasting therapeutic actions of psychedelics, which do not directly involve vision. Therefore, connecting these findings, and psilocybin-induced cortico-cortical circuit alterations more broadly, to therapeutic effects requires follow-up studies. Our results here provide a simple, rapid (<20 minutes with setup), and non-invasive approach for addressing this question in clinical settings while also providing a translational system for investigating cell and circuit-level mechanisms.

One limitation of our study is that the human data come from a retrospective between-subjects design. Although groups were matched on age, sex, and personality metrics, the PSY group had mildly increased health/safety risk taking and greater marijuana usage (Table 1). Importantly, excluding frequent marijuana users did not change the pattern of effects, and most effects identified correlated with time-since psychedelics use. With regard to personality differences, we highlight three subjects in the control group who have used psychedelics in their past (gray points in Figs. 1 and 2); these subjects were not outliers in any major effects. Further, the subsequent mouse experiments helped to further establish a causal relationship between psychedelic use and visual cortical processing. Our second sample of psychedelics users used 5-MeO-DMT, which, like psilocybin and LSD, is a serotonergic psychedelic that activates 5-HT_2A_ receptors, yet 5-MeO-DMT also exhibits strong affinity for 5-HT_1A_ receptors (*66*). Still, 5-MeO-DMT induces similar spine growth in mPFC neurons to psilocybin, and also induces head-twitch responses in mice (*67*). Interestingly, we did not observe as strong correlations with time since use in the 5-MeO-DMT (Supplemental fig. S4D), yet this could be due to the fact that three subjects had also done psilocybin within a week prior to their 5-MeO-DMT dose. Ultimately, future studies should validate our findings by including multiple timepoints from the same individuals, with random assignment and tight control of psilocybin dose.

Another limitation of our study is that our paradigms were not precisely equivalent between these species, although both quantified rapidly emergent (same day, first exposure) predictive suppression of early visual cortical responses (see **Supplemental methods: Paradigm choice**). And while our study focuses on visual processing as a model system, we hypothesize that similar mechanisms of relaxed priors and increased bottom-up signaling may occur across multiple levels of the cortical hierarchy. Future work is needed to understand the connection between sensory cortical processing changes and the therapeutic effects on mood, anxiety, and addiction.

## METHODS

### Humans

All experimental procedures were approved by the Georgia State University Institutional Review Board (IRB) and were carried out in accordance with their guidelines. Participants were recruited through advertisements posted on university bulletin boards, social media platforms, and in local psychedelic community groups. Eligibility criteria included adults aged 18 or older with no history of autism spectrum disorders, bipolar disorder, schizophrenia, current major depressive episodes, or substance abuse, excluding nicotine. Participants filled out a detailed questionnaire to ensure suitability for study participation. Participants provided informed consent and were compensated $30 for their participation in the study.

The study cohort comprised 47 individuals, divided into a control group and a psychedelic group. The control group consisted of 16 participants (6 male, 10 female), who had no recent (within 30 days) history of psychedelic use. The psychedelic group included 31 participants who had used psychedelics within 30 days prior to their participation in this study. This group was further subdivided based on the type of psychedelic used. The PSY subgroup consisted of 16 participants, with 15 taking mushrooms (7 male, 8 female) and 1 (male) taking LSD. Another subgroup comprised 12 participants (5 male, 7 female) who inhaled Incilius alvarius toad toxin. 5-MeO-DMT is considered to be the primary compound present that is responsible for the toxin’s psychoactive effects.

Dosage information for psilocybin, LSD, or 5-MeO-DMT was unattainable, as potency of the drugs used could not be ascertained directly, but participants were pre-screened to exclude those who only took sub-perceptual microdoses. All doses reported were judged as moderate to high. Additionally, within the psychedelic group, 3 out of 16 participants identified as non-binary. Due to the small number of participants in this category, statistical analysis on gender was not feasible. Consequently, gender was excluded from the comparative analysis, although effects of sex and age on all major group differences were analyzed (Supplemental Fig. S5).

#### Saccadic Prediction Task

The saccadic prediction task was conducted in a darkened room where participants were seated 100cm away from an LCD monitor (19-27 inches, 60Hz refresh rate). This task involved visual stimuli in the form of differently colored stars (2.46 degrees of visual angle in diameter), each appearing at one of eight designated spots 8 degrees from fixation on a circular layout. Participants were instructed to initially fixate on a central cross displayed on the screen for 1000ms, followed by shifting their gaze to the star, which appeared for the subsequent 1000ms at one of the eight spots. Each of the 142 trials per run was classified into three types, differentiated by the color of the star: ’expected’ trials featured a green or purple star appearing in a predictable sequence at the same location; ’deviant-update’ trials involved a green or purple star appearing unexpectedly at a new location, signaling participants to update their expectations; and ’deviant-no-update’ trials, indicated by an orange star, appeared unexpectedly but did not require a change in expectation for future star locations. This color-coding scheme was not described to participants prior to the task, but allowed them to learn the significance of each color and to anticipate whether an update in expectation was necessary. Participants completed two consecutive runs for a total duration of nine minutes and forty-seven seconds.

#### EEG recordings and data processing

EEG recordings and data processing were conducted using a 32-channel BioSemi ActiveTwo EEG system. Participants were fitted with a BioSemi 32- channel cap, arranged according to the 10-20 system for electrode placement. EEG data were recorded with a sampling rate of 500 Hz, using a band-pass filter of 0.16 Hz and a lowpass filter at 200 Hz. Reference electrodes were placed 1 cm to the right of Cz (later average referenced) and the ground electrode was positioned at 1 cm to the left of Cz. Impedance was tested prior to each run (SPT1, SPT2, Control run, Oddball, Oddball flip), and kept below 10kΩ for each electrode.

Data preprocessing was performed using Besa Research software (Gräfelfing, Germany) as previously described (*38*, *45*). Data were average referenced and noisy channels were interpolated (two or fewer per run). Eye-blinks and saccades were identified using Independent Components Analyses (ICA) and removed from the EEG data (*68*). Individual trial data was then isolated for subsequent analysis (-700 to 1500ms pre- vs post-stimulus onset for SPT runs; - 375ms to 875ms pre- to post-stimulus onset for oddball runs). For EEG analyses, SPT trials were excluded if there was no saccade, or if there was a saccade from -300ms to 150ms post-stimulus onset to ensure adequate processing of early stimulus evoked activity. Data was baseline-corrected (-200 to 0 ms pre-stimulus onset) and averaged for plotting event related potentials (Fig. 2A-F), which was done mainly for descriptive purposes. The primary analyses of all human and mouse data were done in the time-frequency domain in Matlab (Mathworks, Natick, MA, USA), using the EEGLAB toolbox (*68*).

Individual trial data for each electrode was converted to the time-frequency domain with a modified morelet wavelet approach with 100 evenly spaced wavelets from 1 to 100 Hz (for SPT) or 2 to 100Hz (for oddball, as stimuli were only 500ms long, activity < 2Hz is difficult to interpret), linearly increasing in length from 1 to 25 cycles per wavelet, applied every 10ms. Stimulus-induced power spectra was computed as decibels relative to the pre-stimulus baseline (-200 to 0ms) for each frequency, timepoint, electrode, trial, condition, and participant, and averaged over trials to yield time-frequency power plots for each electrode, condition, and participant. Comparisons were carried out using traditional frequency bands (delta: 1-4Hz, theta: 5-8Hz (5-10 for mice); alpha: 9-14Hz, beta: 15-34Hz, gamma: 35–100Hz) and on two separate time windows based on maximal time and time-frequency responses in the grand average plots: early (50-300ms) and late (300-600ms). These two windows also map onto the established latencies of typically analysed early (e.g., N100, MMN) vs late (P300) evoked cortical activities (ERPs), respectively. We also separately examined effects of prediction (standard vs all deviants) and updating (deviant update vs deviant non-update). In order to minimize statistical comparisons, tests were carried out after averaging over all electrodes, as task related brain activity was largely scalp-wide (See Fig. 2H). The number of trials used for each trial type was equated across conditions for the SPT (n=40 for standard and deviant locations).

Additionally, intertrial phase locking (phase-locking factor, or PLF) was analyzed for the SPT task in order to help isolate brain activity that was locked to stimulus onset (i.e., not convolved with e.g., response preparation). PLF was calculated by dividing the complex results of the above described wavelet analysis by their absolute values for each electrode, timepoint, frequency, participant, trial, and condition. The resulting values were averaged across trials, with the absolute value of this result yielding the PLF or R-statistic, which is bound between 0 and 1 (1 indicating perfect phase alignment across trials). In non-phaselocked (or phase-randomized) data, R-values vary depending on the number of trials involved in the calculation and can be baseline corrected using previously described mathematical approaches(*45*, *69*), as they were done in this study.

#### Saccade scoring

Eye movements were measured using two additional facial electrodes (BioSemi) placed 1cm below and beside the outer canthus of the left eye (Supplemental Fig. S1A). All saccades were manually scored using routines written in the host lab. Scoring focused on the onset of the first saccade after the onset of each stimulus and used a combination of each electrode and the average of their absolute value. Each run and subject was scored by at least 2 scorers, blinded to group membership or trial type. In the case where discrepancies were present (lower than .85 inter-rater reliability for a given subject, across all trials; or greater than 20ms difference between trial), a third scorer was included. Scores were averaged between the two closest estimates. Trials without clear saccades in the 800ms after stimulus onset, or with saccades present in the 300ms prior to trial onset were not scored and were excluded from EEG analysis as well. In the end, all subjects had greater than 75% of trials included with greater than .85 inter-rater reliability. The proportion of trials excluded did not differ between groups, t(29)=1.46,p=.16). Saccades with latencies >500ms were rare (<3.12% overall) and did not differ between groups (F^group^(1,29)=0.5; p=.48) or show a group by stimulus location interaction (F^interaction^ (1,29)=0.14; p=.71). As such slow responses could reflect task disengagement, we excluded them from subsequent analyses. Saccades with latencies greater than 75ms and less than 147ms were considered express saccades based on i) past literature(*36*), ii) neuroanatomy(*37*), iii) group-level histograms (Fig. 1B), and iv) a curve fitting procedure using Matlab. In (iv), we fit three gaussians to the overall distribution across all PSY and CNT subjects. The main (largest) gaussian captured 77.9% of responses and had a mean of 271.2 ms and a standard deviation of 58ms. The second population captured express and anticipatory saccades as 9% of the total, and had a mean of 75.7ms and a standard deviation of 42.5ms. In this population, the 95%ile upper cutoff was 146.8ms, and so we considered responses shorter than that, but higher than the typical “anticipatory” threshold to be “express” responses. The third population had a large spread from 140ms (5%ile) to 642.3 (95%ile) and was platykurtic, with a mean of 391.6ms and a standard deviation of 150ms.

### Mice

#### Animals

All animals were housed at a 12/12 light/dark cycle with food and water available *ad libitum* at the Nathan Kline Institute (NKI) animal facilities. All experiments were performed under the approval of the institutional animal care and use committee (IACUC) at NKI. Adult C57BL/6J mice, n = 25, age range from P87 to P183 (from Jackson Laboratories) were used. Both female (n=11) and male mice (n=14) were included.

#### Surgeries

For stereotaxic surgery, animals were anesthetized using 3% isoflurane, maintained at 1-2% isoflurane, and received pre and post care medication appropriately. For local field potential (LFP) recordings, small 0.2 mm craniotomies were performed at mouse V1 (coordinates from bregma: X =-2mm, Y = -2.92mm) and ACa (X=0.3 mm, Y=0.6 mm) and bipolar electrodes were implanted in each region, targeting layer 2/3 (200µm below the pial surface), and grounded at an adjacent skull surface. Prior to insertion, electrodes were submerged in DiI dye for post-hoc anatomical validation. During the same surgery session, a titanium head-plate was secured at the mouse head to allow for their fixation to the imaging apparatus.

For spine analysis, AAVrg-hSyn-Cre (Addgene #105553, 2.5x10^13^ GC/ml) was injected into V1, mixture of AAV1-Flex-CAG-EGFP (Addgene #51502, 2.5x10^13^ GC/ml) and AAV1-Flex-hSyn-hm4di-mCherry (Addgene #44362, 2.4x10^13^ GC/ml) at a ratio 1:19 into ACa. Following coordinates were used. V1: 200 nl (from bregma: X =-2.2mm, Y = -2.9mm, Depth from the dorsal surface: 0.5mm), ACa: 200 nl/sites, 3 sites (X: 0.3mm, Y: 0mm, Depth 0.6mm, X: 0.4 mm, Y: 1.0 mm, Depth 1.1 mm, X: 0.4mm, Y: 2mm, Depth 1.0 mm).

#### Drug administration and observation

Saline or Psilocybin (Usona Institute, 1mg/kg, intraperitoneally) was injected after the first LFP recordings. Mice were then returned to their home cages. In a subset of mice, we recorded their movements 15 minutes after dosing in the home cage using an iPhone 16 (apple) at 60 frames per second. Head-twich responses during these 30 minute videos were then scored manually, blind to drug group.

#### Global-local Oddball Paradigm

Visual stimuli were presented on a computer monitor (19-inch diameter, 60Hz refresh rate) at a 45-degree angle from the animal axis, approximately 15cm from the mouse’s eyes, using the Psychophysics Toolbox on MATLAB (Mathworks) while the mouse was head-fixed to table. Visual stimuli consisted of black and white, full-field square-wave gratings at approximately 0.08 cycles per degree, drifting at 2 cycles per second, each presented for 500ms and separated from one another by 450-550ms of black screen.

For the global/local paradigm, short sequences of five stimuli of 500ms in duration, with an inter-stimulus interval of 500-550ms of gray screen, were presented, followed by 5000-5500 ms of gray screen (inter-sequence interval). Initially, mice viewed a “training set”, which included 70 sequences of AAAAB where A was 0° and B was 90°. After this, mice rested for 2-3 minutes, and then viewed a “test set”, wherein AAAAB sequences were primarily presented, but AAAAA (global deviant, local redundant) and AAAAC (global deviant, local deviant) catch trials were presented one in 6 presentations (C was always 45 °). Then, after another brief rest, a control run was presented, wherein random orientations (e.g., CDEAB) were presented in the same 5 stimulus sequences (500ms duration, 500-550ms ISI, 5000-5500 ms inter-sequence interval). Random orientations included degree angles: 22.5°, 45°, 67.5°, 90°, 112.5°, 135°, 157.5°, and 180°.

#### Standard Oddball Paradigm

Prior to the global-local runs, mice viewed a standard oddball paradigm, consisting of 4 oddball runs and two many-standards controls. The same stimulus characteristics, duration, and ISIs were used, but stimuli were presented in continuous trains of 240 trials (≈4 minutes). The many-standards control sequence was composed of eight orientations that were presented in random order. The oddball sequence consisted of a repetitive sequence of one stimulus (“redundant”, either 0°, or 90° degree angles, presented 87.5% of the time), randomly interrupted by a stimulus of a different orientation (“deviant”, 90°, or 0° degree angles, presented 12.5% of the time). In a subsequent sequence, the redundant stimulus was “flipped” to become the deviant, and vice versa (“oddball flip”); thus, we can assess responses to the stimulus context (i.e., in what context a stimulus is shown) rather than stimulus features (i.e., what orientation a stimulus is). Oddball runs started with 30-35 repeats of the redundant orientation. All mice completed two rounds of the oddball-oddballflip-control paradigm, testing angles 0°, 90°, 45°, and 135° as deviant, redundant, and control.

#### LFP recordings and data processing

Head-fixation and visual stimulation habituation was performed seven to ten days following the surgery, in increments of 5-minute sessions each day (5 minutes for day one, 10 minutes for day 2, and 15 minutes for day 3), in which the visual stimuli were presented only once (many standards control run). This was done to acclimate the mouse to head-fixation and to reduce movement during recordings. Recordings took place two weeks after post-surgery.

Mice were head-fixed to the recording apparatus and free to move on a manual treadmill during recordings. Mice were monitored by the experimenter to ensure that they were awake and not in clear distress or discomfort during all recordings. Insulated cables were connected to the implanted electrodes and plugged into a differential amplifier (Warner instruments, DP-304A, high-pass: 0 Hz, low-pass: 500 Hz, gain: 1K, Holliston, MA, USA). Amplified signals were passed through a 60 Hz noise cancellation machine (Digitimer, D400, Mains Noise Eliminator, Letchworth Garden City, UK), which, instead of filtering creates an adaptive subtraction of repeating signals which avoids phase delays or other forms of waveform distortion that can result from standard bandstop filters. Electrophysiological activity was recorded using the Prairie View software. Locomotion was recorded with a rotary encoder embedded in the treadmill and connected to the computer as an analogue input. Although locomotion can alter the gain of visually evoked responses (*70*), at least two previous studies (*16*, *71*) have shown that the visual oddball paradigm does not evoke reliable changes in locomotion or other facial motor variables (e.g., whisking) in mice. We found that locomotion was present on <10% of trials for all mice, so trials (fig 3, S6,7) and segments (fig 4, S9) during locomotion were excluded from further analysis, as was done previously (*16*, *71*). This step did not change results or conclusions.

#### Local field potential signal processing and analysis

Trials with excessive signal (>≈5 std devs) in either V1 and ACa were manually excluded (between 0 and 10 for each mouse). Analyses focused on presentations of the same stimulus (0, 45, 90, or 135 degrees) in the neutral vs deviant context (control vs oddball) in both paradigms (standard vs global-local). The number of trials included was equated across conditions and paradigms (between 8 and 12). As such, analyses in the standard oddball were limited to the orientation that evoked the strongest response averaged across contexts (control and deviant/oddball). Only one orientation was presented as the deviant during the global-local (90 or 0 degrees), and we focused on the last 10-12 presentations to maximize estimates predictive suppression(*15*). During the global-local-control (i.e., when orientations were presented randomly in 5 stimulus bursts), we focused on trials in which that orientation was a) appearing in the 4th or 5th slot and b) not preceded by an orientation closer than 45 degrees of angle difference, in order to minimize the short term impacts of stimulus specific adaptation (*42*) and to better match the global-local-oddball condition.

Ongoing data were converted to the time-frequency domain with the same approach described above. Statistical analyses focused on comparisons between neutral and deviant stimulus induced power from delta to theta frequency bands, in the 50ms to 300ms period post stimulus onset to best match the time-frequency regions of interest from the human study. Theta in rodents is notably higher than in humans, so instead of 5-8Hz, we extended it to 10Hz.

For analyzing effective connectivity between ACa and V1, we carried out a non-parametric Granger causality-based approach in the time-frequency domain (*16*, *72*) as implemented in the Fieldtrip toolbox(*73*). This analysis examined lagged spectral covariance between regions for each direction (ACa to future-V1 vs V1 to future-ACa) for each frequency 1-40Hz. Data was binned in 1000ms segments throughout the continues recordings of the global-local oddball and control runs. We focused on a time-lag of 11 to 15 ms based on similar studies of long-range brain connectivity in the mouse(*16*, *74*). The resulting value is a Granger coefficient (GC) reflecting the spectral covariance in between the signal from one region and the lagged signal from the other region.. Plots focused on 1 to 40Hz, as higher frequencies showed small values across mice. Analyses focused on 1 to 5Hz, as this is where the ACa to V1 granger causality was maximal across mice in the top-down direction (Fig. 3; S8). We split data into stimulus and inter-stimulus interval periods, and binned data in 1000ms bins at the start, middle, and end of each period, and then averaged over trials. Granger causality in the top-down direction was ostensibly stronger during the stimulus period across all mice, but this was not statistically significant (F^directionXperiod^(1,209)=2.02, p=.15). Focusing analyses only on the stimulus period, group differences remained (F^directionXgroup^(1,95)=7.57, p=.007).

#### Histology

Mice were deeply anesthetized via urethane and 3% isoflurane and then perfused transcardially with PBS followed by 4% paraformaldehyde (PFA)/PBS. Heads with electrodes and headplates attached were post-fixed in 4% PFA/PBS for two overnights. Four mice (2 of each group) required additional immunostaining. After isolation, brains were cryoprotected with 30% sucrose/PBS for one to two overnights and then embedded in Tissue-Tek OCT compound (Sakura) and cryosectioned (50 µm). For spine analysis, sections were stained with anti-GFP (ThermoFisher, A11122, 1/1000), anti-rabbit IgG-Alexa488 (Jackson Immuno, #711-545-152, 1/500), DAPI (ThermoFisher, D1306, 1 µg/ml) in PBST (0.1% Tween20) and mounted using an antifade medium (VectorLabs, H-1700-2).Sections were scanned using LSM880 (Zeiss) confocal microscopy. Images were analyzed using FIJI and RESPAN with a pre-trained model (Model 1A). Staining method and batch was included as a fixed covariate to account for procedural differences across a subset of animals.

### Statistical analyses

All statistics were carried out in Matlab. In the human data, all measures and effects reported in figures 1-2 and supplemental figures S1-S5, S8 came from the same 47 individuals, from either the SPT task (2 runs combined) or the oddball paradigm (S8). For saccade analyses, we used Komologrov-Smirnov tests on mean-centered saccade distributions, collapsed over all participants in each group, to test for differences in latency distributions. We used mixed ANOVAs on saccade latency and proportion of express saccades, with GROUP as a between-subjects factor (CNT, PSY or 5-MeO-DMT) and LOCATION as a within-subjects factor (standard, deviant). For SPT task EEG activity, we used mixed ANOVAs on early and late time period power and PLF, with GROUP as a between-subjects factor (CNT, PSY or 5-MeO-DMT) and LOCATION as a within-subjects factor (standard, deviant). For both saccades and EEG data during the SPT, we focused analyses on the second through fourth standards in the sequence in order to equate the number of trials across conditions and avoid updating effects (see Fig. 1F,G). Analyses were carried out on the average across all electrodes to minimize statistical tests, and separately for delta, theta, alpha, beta, and gamma bands. Significant interactions were followed up with paired or two-sample t-tests with one-tailed significance, as greater activity to the deviant vs the standard was the clear a priori hypothesis.

In the mouse data, in all cases (stimulus induced power, Granger coefficients, spine metrics), we carried out linear mixed effects (LME) models with mouse as a random effects variable, and sex included as a covariate. Functional data reported in figures 3-4 and supplemental figures S6, S7, and S9 came from the same 19 mice, recorded during oddball paradigms. Structural data came from an additional 12 mice. To compare visual processing in mice, we focused on the post dose time point (24 hours) for the main analyses. We averaged activity across trials (see above) within the time bin (50-300ms) and frequency band (delta-theta) that differed in the human part of the study, in order to minimize statistical comparisons. We included fixed-effects terms for treatment (psilocybin vs saline), paradigm (global-local vs standard oddball), and context (neutral vs deviant), in addition to all second and third-order interactions. The model was restricted to the three primary interaction terms of interest to preserve statistical power and avoid overfitting. Sex was included as a main effect covariate rather than an interaction term, as the data were insufficiently powered to reliably estimate higher-order interactions. Ostensibly, effects were present regardless of sex. The variation for repeated measures within mouse was accounted for by including mouse as a random effects term. Significant interactions were followed up with two-sample t-tests with two-tailed significance during the global-local oddball (as responses could be augmented or suppressed, depending on whether top-down suppression is intact) and one-tailed significance during the standard oddball (as numerous past studies have all observed the same increase in delta-theta power to deviant stimuli). For the Granger causality analysis, averaged across lags (11 to 12ms) and runs (oddball and control) and low frequencies (2-4Hz, based on overall average curve, figure 3) for each mouse, direction (top-down vs bottom-up), bin (beginning, middle, end of segment), and treatment (SAL vs PSL). Again, the model was restricted to the three primary interaction terms of interest (bin, direction, treatment) to preserve statistical power and avoid overfitting, while sex was included as a fixed effect variable and mouse as a random effects term.

To analyse spine density, single measures for each dendritic segment were analysed in an LME with treatment (SAL vs PSL) and sex as fixed effects terms, along with a sex by treatment interaction. We also included batch (three separate batches with equal numbers from each group) as a fixed effect variable and mouse as a random effects term. The same model was used for spine volume.

## Data availability

The processed EEG, EOG, and LFP, and structural data (.tif files) data will be made available upon publication. Raw data will be made available upon reasonable request.

## Code availability

Data analysis was carried out with Besa Research software (Gräfelfing, Germany), using FUJI, or in Matlab using the Fieldtrip toolbox, EEGlab toolbox, and custom code, which will be made publicly available upon publication.

## Acknowledgments

This work was funded by the National Eye Institute (R01EY033950 to J.P.H.) and the Brain and Behavior Research Foundation (YI30149 to J.P.H.). We also acknowledge Usona Institute for providing Psilocybin through their Investigational Drugs and Materials Supply Program.

## References

1. G. M. Goodwin, S. T. Aaronson, O. Alvarez, P. C. Arden, A. Baker, J. C. Bennett, C. Bird, R. E. Blom, C. Brennan, D. Brusch, L. Burke, K. Campbell-Coker, R. Carhart-Harris, J. Cattell, A. Daniel, C. DeBattista, B. W. Dunlop, K. Eisen, D. Feifel, M. Forbes, H. M. Haumann, D. J. Hellerstein, A. I. Hoppe, M. I. Husain, L. A. Jelen, J. Kamphuis, J. Kawasaki, J. R. Kelly, R. E. Key, R. Kishon, S. Knatz Peck, G. Knight, M. H. B. Koolen, M. Lean, R. W. Licht, J. L. Maples-Keller, J. Mars, L. Marwood, M. C. McElhiney, T. L. Miller, A. Mirow, S. Mistry, T. Mletzko-Crowe, L. N. Modlin, R. E. Nielsen, E. M. Nielson, S. R. Offerhaus, V. O’Keane, T. Páleníček, D. Printz, M. C. Rademaker, A. van Reemst, F. Reinholdt, D. Repantis, J. Rucker, S. Rudow, S. Ruffell, A. J. Rush, R. A. Schoevers, M. Seynaeve, S. Shao, J. C. Soares, M. Somers, S. C. Stansfield, D. Sterling, A. Strockis, J. Tsai, L. Visser, M. Wahba, S. Williams, A. H. Young, P. Ywema, S. Zisook, E. Malievskaia, Single-dose psilocybin for a treatment-resistant episode of major depression. N. Engl. J. Med. 387, 1637–1648 (2022).

2. D. Nutt, R. Carhart-Harris, The current status of psychedelics in psychiatry. JAMA Psychiatry 78, 121–122 (2021).

3. B. Kelmendi, A. P. Kaye, C. Pittenger, A. C. Kwan, Psychedelics. Curr. Biol. 32, R63–R67 (2022).

4. H. M. Dourron, C. D. Nichols, O. Simonsson, M. Bradley, R. Carhart-Harris, P. S. Hendricks, 5-MeO-DMT: An atypical psychedelic with unique pharmacology, phenomenology & risk? Psychopharmacology (Berl.) 242, 1457–1479 (2025).

5. K. Alper, J. Cange, R. Sah, D. Schreiber-Gregory, H. Sershen, K. Y. Vinod, Psilocybin sex-dependently reduces alcohol consumption in C57BL/6J mice. Front. Pharmacol. 13, 1074633 (2022).

6. N. K. Savalia, L.-X. Shao, A. C. Kwan, A dendrite-focused framework for understanding the actions of ketamine and psychedelics. Trends Neurosci. 44, 260–275 (2021).

7. L.-X. Shao, C. Liao, I. Gregg, P. A. Davoudian, N. K. Savalia, K. Delagarza, A. C. Kwan, Psilocybin induces rapid and persistent growth of dendritic spines in frontal cortex in vivo. Neuron 109, 2535–2544.e4 (2021).

8. C. Ly, A. C. Greb, L. P. Cameron, J. M. Wong, E. V. Barragan, P. C. Wilson, K. F. Burbach, S. Soltanzadeh Zarandi, A. Sood, M. R. Paddy, W. C. Duim, M. Y. Dennis, A. K. McAllister, K. M. Ori-McKenney, J. A. Gray, D. E. Olson, Psychedelics promote structural and functional neural plasticity. Cell Rep. 23, 3170–3182 (2018).

9. Q. Jiang, L.-X. Shao, S. Yao, N. K. Savalia, A. D. Gilbert, P. A. Davoudian, J. D. Nothnagel, G. Tian, T. S. Hung, H. M. Lai, K. T. Beier, H. Zeng, A. C. Kwan, Psilocybin triggers an activity-dependent rewiring of large-scale cortical networks. Cell 189, 659–675.e22 (2026).

10. R. Moliner, M. Girych, C. A. Brunello, V. Kovaleva, C. Biojone, G. Enkavi, L. Antenucci, E. F. Kot, S. A. Goncharuk, K. Kaurinkoski, M. Kuutti, S. M. Fred, L. V. Elsilä, S. Sakson, C. Cannarozzo, C. R. A. F. Diniz, N. Seiffert, A. Rubiolo, H. Haapaniemi, E. Meshi, E. Nagaeva, T. Öhman, T. Róg, E. Kankuri, M. Vilar, M. Varjosalo, E. R. Korpi, P. Permi, K. S. Mineev, M. Saarma, I. Vattulainen, P. C. Casarotto, E. Castrén, Psychedelics promote plasticity by directly binding to BDNF receptor TrkB. Nat. Neurosci. 26, 1032–1041 (2023).

11. L.-X. Shao, C. Liao, P. A. Davoudian, N. K. Savalia, Q. Jiang, C. Wojtasiewicz, D. Tan, J. D. Nothnagel, R.-J. Liu, S. C. Woodburn, O. M. Bilash, H. Kim, A. Che, A. C. Kwan, Psilocybin’s lasting action requires pyramidal cell types and 5-HT2A receptors. Nature 642, 411–420 (2025).

12. A. B. Ozols, J. Wei, J. M. Campbell, C. Hu, S. Qiu, A. L. Gallitano, Activity of prefrontal cortex serotonin 2A receptor expressing neurons is necessary for the head-twitch response of mice to psychedelic drug DOI in a sex-dependent manner, bioRxivorg (2024). 10.1101/2024.05.21.595211.

13. J. S. Siegel, C. Liston, G. E. Nicol, R. L. Carhart-Harris, M. P. Bogenschutz, The science of psychedelic medicine. Nat. Med. 32, 449–462 (2026).

14. A. Barstowe, P. J. Kajonius, Masking influences: A systematic review of placebo control and masking in psychedelic studies. J. Psychoactive Drugs 58, 7–17 (2026).

15. D. S. Peterka, F. Imai, J. M. Ross, G. Bastos, M. Hornick, L. Rachmany, C. G. Gallimore, A. Hockley, J. P. Hamm, Global context rapidly shapes sensory responses in V1, bioRxivorg (2026). 10.64898/2026.01.07.698143.

16. G. Bastos, J. T. Holmes, J. M. Ross, A. M. Rader, C. G. Gallimore, J. A. Wargo, D. S. Peterka, J. P. Hamm, Top-down input modulates visual context processing through an interneuron-specific circuit. Cell Rep. 42, 113133 (2023).

17. V. S. Raghavan, J. Madsen, M. Nentwich, M. Leszczynski, A. Falchier, S. Bickel, B. E. Russ, L. C. Parra, During natural vision, semantic novelty modulates fixation-related processing in primate cortex, bioRxivorg (2026)p. 2026.03.18.712708.

18. M. Solyga, M. Zelechowski, G. B. Keller, Visuomotor mismatch EEG responses over occipital cortex of freely moving human subjects, bioRxiv (2025). 10.1101/2025.08.14.670295.

19. E. Sennesh, J. A. Westerberg, J. Spencer-Smith, A. Bastos, Ubiquitous predictive processing in the spectral domain of sensory cortex, eLife (2025). 10.7554/elife.109053.1.

20. P. A. Davoudian, L.-X. Shao, A. C. Kwan, Shared and distinct brain regions targeted for immediate early gene expression by ketamine and psilocybin. ACS Chem. Neurosci. 14, 468–480 (2023).

21. K. Friston, A theory of cortical responses. Philos. Trans. R. Soc. Lond. B Biol. Sci. 360, 815–836 (2005).

22. A. M. Bastos, W. M. Usrey, R. A. Adams, G. R. Mangun, P. Fries, K. J. Friston, Canonical microcircuits for predictive coding. Neuron 76, 695–711 (2012).

23. R. L. Carhart-Harris, K. J. Friston, REBUS and the anarchic brain: Toward a unified model of the brain action of psychedelics. Pharmacol. Rev. 71, 316–344 (2019).

24. S. Zhang, M. Xu, T. Kamigaki, J. P. Hoang Do, W.-C. Chang, S. Jenvay, K. Miyamichi, L. Luo, Y. Dan, Selective attention. Long-range and local circuits for top-down modulation of visual cortex processing. Science 345, 660–665 (2014).

25. C. M. Niell, M. P. Stryker, Highly selective receptive fields in mouse visual cortex. J. Neurosci. 28, 7520–7536 (2008).

26. J. P. Hamm, Y. Shymkiv, S. Han, W. Yang, R. Yuste, Cortical ensembles selective for context. Proc. Natl. Acad. Sci. U. S. A. 118, e2026179118 (2021).

27. A. Bravermanová, M. Viktorinová, F. Tylš, T. Novák, R. Androvičová, J. Korčák, J. Horáček, M. Balíková, I. Griškova-Bulanova, D. Danielová, P. Vlček, P. Mohr, M. Brunovský, V. Koudelka, T. Páleníček, Psilocybin disrupts sensory and higher order cognitive processing but not pre-attentive cognitive processing-study on P300 and mismatch negativity in healthy volunteers. Psychopharmacology (Berl.) 235, 491–503 (2018).

28. P. Duerler, S. Brem, G. Fraga-González, T. Neef, M. Allen, P. Zeidman, P. Stämpfli, F. X. Vollenweider, K. H. Preller, Psilocybin induces aberrant prediction error processing of tactile mismatch responses-A simultaneous EEG-FMRI study. Cereb. Cortex 32, 186–196 (2021).

29. D. Umbricht, F. X. Vollenweider, L. Schmid, C. Grübel, A. Skrabo, T. Huber, R. Koller, Effects of the 5-HT2A agonist psilocybin on mismatch negativity generation and AX-continuous performance task: implications for the neuropharmacology of cognitive deficits in schizophrenia. Neuropsychopharmacology 28, 170–181 (2003).

30. C. H. Murray, I. Tare, C. M. Perry, M. Malina, R. Lee, H. de Wit, Low doses of LSD reduce broadband oscillatory power and modulate event-related potentials in healthy adults. Psychopharmacology (Berl.) 239, 1735–1747 (2022).

31. A. M. Michaiel, P. R. L. Parker, C. M. Niell, A hallucinogenic serotonin-2A receptor agonist reduces visual response gain and alters temporal dynamics in mouse V1. Cell Rep. 26, 3475–3483.e4 (2019).

32. T. Lyons, M. Spriggs, L. Kerkelä, F. E. Rosas, L. Roseman, P. A. M. Mediano, C. Timmermann, L. Oestreich, B. A. Pagni, R. J. Zeifman, A. Hampshire, W. Trender, H. M. Douglass, M. Girn, K. Godfrey, H. Kettner, F. Sharif, L. Espasiano, A. Gazzaley, M. B. Wall, D. Erritzoe, D. J. Nutt, R. L. Carhart-Harris, Human brain changes after first psilocybin use. Nat. Commun. 17 (2026).

33. McCrae, Costa, Personality in Adulthood (Guilford Publications, New York, NY, 2003).

34. J. X. O’Reilly, U. Schüffelgen, S. F. Cuell, T. E. J. Behrens, R. B. Mars, M. F. S. Rushworth, Dissociable effects of surprise and model update in parietal and anterior cingulate cortex. Proc. Natl. Acad. Sci. U. S. A. 110, E3660–9 (2013).

35. E. Kayhan, S. Hunnius, J. X. O’Reilly, H. Bekkering, Infants differentially update their internal models of a dynamic environment. Cognition 186, 139–146 (2019).

36. B. Fischer, H. Weber, M. Biscaldi, F. Aiple, P. Otto, V. Stuhr, Separate populations of visually guided saccades in humans: reaction times and amplitudes. Exp. Brain Res. 92, 528–541 (1993).

37. J. E. McDowell, K. A. Dyckman, B. P. Austin, B. A. Clementz, Neurophysiology and neuroanatomy of reflexive and volitional saccades: evidence from studies of humans. Brain Cogn. 68, 255–270 (2008).

38. J. P. Hamm, K. A. Dyckman, L. E. Ethridge, J. E. McDowell, B. A. Clementz, Preparatory activations across a distributed cortical network determine production of express saccades in humans. The Journal of Neuroscience 30, 7350–7357 (2010).

39. B. A. Clementz, The ability to produce express saccades as a function of gap interval among schizophrenia patients. Exp. Brain Res. 111, 121–130 (1996).

40. B. Fischer, E. Ramsperger, Human express saccades: effects of randomization and daily practice. Exp. Brain Res. 64, 569–578 (1986).

41. R. M. Müri, S. Rivaud, B. Gaymard, C. J. Ploner, A. I. Vermersch, C. W. Hess, C. Pierrot-Deseilligny, Role of the prefrontal cortex in the control of express saccades. A transcranial magnetic stimulation study. Neuropsychologia 37, 199–206 (1999).

42. J. P. Hamm, R. Yuste, Somatostatin interneurons control a key component of mismatch negativity in mouse visual cortex. Cell Rep. 16, 597–604 (2016).

43. D. C. Javitt, M. Lee, J. T. Kantrowitz, A. Martinez, Mismatch negativity as a biomarker of theta band oscillatory dysfunction in schizophrenia. Schizophr. Res. 191, 51–60 (2018).

44. B. J. Roach, D. H. Mathalon, Event-related EEG time-frequency analysis: an overview of measures and an analysis of early gamma band phase locking in schizophrenia. Schizophr. Bull. 34, 907–926 (2008).

45. J. P. Hamm, K. A. Dyckman, J. E. McDowell, B. A. Clementz, Pre-Cue Fronto-Occipital Alpha Phase and Distributed Cortical Oscillations Predict Failures of Cognitive Control. The Journal of Neuroscience 32, 7034–7041 (2012).

46. W. J. Cubala, M. Bajbouj, M. Bauer, B. T. Baune, N. Cardoner, F. Devlin, K. Doolin, R. M. Dueñas Herrero, M. Elices, A. Feeney, M. Galuszko-Wegielnik, K. Jakuszkowiak-Wojten, L. Janu, J. R. Kelly, K. Ledden, R. Maclsaac, S. Madero, S. J. McInerney, A. L. Montejo, A. Nawka, T. Pálenícek, V. Pérez Solà, J. G. Ramaekers, A. Reif, P. Ritter, F. Ryan, C. B. Svendsen, C. Sweeney, T. H. Terwey, M. H. Trivedi, V. Valcheva, E. Vieta, M. E. Thase, GH001 vs placebo in patients with treatment-resistant depression: A randomized clinical trial. JAMA Psychiatry, doi: 10.1001/jamapsychiatry.2026.0096 (2026).

47. L. Harms, P. T. Michie, R. Näätänen, Criteria for determining whether mismatch responses exist in animal models: Focus on rodents. Biol. Psychol. 116, 28–35 (2016).

48. J. A. Westerberg, Y. S. Xiong, E. Sennesh, H. Nejat, D. Ricci, S. Durand, B. Hardcastle, H. Cabasco, H. Belski, A. Bawany, R. Gillis, H. Loeffler, C. R. Peene, W. Han, K. Nguyen, V. Ha, T. Johnson, C. Grasso, A. Young, J. Swapp, B. Ouellette, S. Caldejon, A. Williford, P. A. Groblewski, S. R. Olsen, C. Kiselycznyk, C. Koch, J. A. Lecoq, A. Maier, A. M. Bastos, Hierarchical substrates of prediction in visual cortical spiking, bioRxivorg (2025). 10.1101/2024.10.02.616378.

49. S. Furutachi, A. D. Franklin, A. M. Aldea, T. D. Mrsic-Flogel, S. B. Hofer, Cooperative thalamocortical circuit mechanism for sensory prediction errors. Nature 633, 398–406 (2024).

50. R. Jordan, G. B. Keller, Opposing influence of top-down and bottom-up input on excitatory layer 2/3 neurons in mouse primary visual cortex. Neuron 108, 1194–1206.e5 (2020).

51. R. P. Rao, D. H. Ballard, Predictive coding in the visual cortex: a functional interpretation of some extra-classical receptive-field effects. Nat. Neurosci. 2, 79–87 (1999).

52. S. B. Garcia, A. P. Schlotter, D. Pereira, A. J. Recupero, F. Polleux, L. A. Hammond, RESPAN: A deep learning pipeline for accurate and automated restoration, segmentation, and quantification of dendritic spines, bioRxivorg (2025). 10.1101/2024.06.06.597812.

53. A. Hockley, L. H. Bohórquez, M. S. Malmierca, Top-down prediction signals from the medial prefrontal cortex govern auditory cortex prediction errors. Cell Rep. 44, 115538 (2025).

54. R. J. Zeifman, M. J. Spriggs, H. Kettner, T. Lyons, F. E. Rosas, P. A. M. Mediano, D. Erritzoe, R. L. Carhart-Harris, From relaxed beliefs under psychedelics (REBUS) to revised beliefs after psychedelics (REBAS). Sci. Rep. 15, 3651 (2025).

55. 55. I. Aizenbud, N. Audette, R. Auksztulewicz, K. Basiński, A. M. Bastos, M. Berry, A. Canales-Johnson, H. Choi, C. Clopath, U. Cohen, R. P. Costa, R. De Filippo, R. Doronin, S. P. Errington, J. P. Gavornik, C. J. Gillon, A. Granier, J. P. Hamm, L. Hertäg, H. Kennedy, S. Kumar, A. Ladd, H. Ladret, J. A. Lecoq, A. Maier, P. McCarthy, J. Mei, J. Mejias, F. Mikulasch, N. Mudrik, F. Najafi, K. Nejad, H. Nejat, K. Oweiss, M. A. Petrovici, V. Priesemann, L. Rudelt, S. Ruediger, S. Russo, A. Salatiello, W. Senn, E. Sennesh, S. Sima, C. Uran, A. Vasilevskaya, J. Vezoli, M. Vinck, J. A. Westerberg, K. Wilmes, Y. S. Xiong, Neural mechanisms of predictive processing: a collaborative community experiment through the OpenScope program, arXiv [q-bio.NC] (2025). http://arxiv.org/abs/2504.09614.

56. A. Fiser, D. Mahringer, H. K. Oyibo, A. V. Petersen, M. Leinweber, G. B. Keller, Experience-dependent spatial expectations in mouse visual cortex. Nat. Neurosci. 19, 1658–1664 (2016).

57. J. M. Ross, J. P. Hamm, An expanding repertoire of circuit mechanisms for visual prediction errors. Trends Neurosci., doi: 10.1016/j.tins.2024.10.007 (2024).

58. L. J. Mertens, M. B. Wall, L. Roseman, L. Demetriou, D. J. Nutt, R. L. Carhart-Harris, Therapeutic mechanisms of psilocybin: Changes in amygdala and prefrontal functional connectivity during emotional processing after psilocybin for treatment-resistant depression. J. Psychopharmacol. 34, 167–180 (2020).

59. J. S. Siegel, S. Subramanian, D. Perry, B. P. Kay, E. M. Gordon, T. O. Laumann, T. R. Reneau, N. V. Metcalf, R. V. Chacko, C. Gratton, C. Horan, S. R. Krimmel, J. S. Shimony, J. A. Schweiger, D. F. Wong, D. A. Bender, K. M. Scheidter, F. I. Whiting, J. A. Padawer-Curry, R. T. Shinohara, Y. Chen, J. Moser, E. Yacoub, S. M. Nelson, L. Vizioli, D. A. Fair, E. J. Lenze, R. Carhart-Harris, C. L. Raison, M. E. Raichle, A. Z. Snyder, G. E. Nicol, N. U. F. Dosenbach, Psilocybin desynchronizes the human brain. Nature 632, 131–138 (2024).

60. J. P. Hamm, “Analyzing functional neuronal ensembles in a between-subjects paradigm” in Neuromethods (Springer US, New York, NY, 2025)*Neuromethods*, pp. 219–234.

61. J. P. Hamm, Y. Shymkiv, J. Mukai, J. A. Gogos, R. Yuste, Aberrant cortical ensembles and schizophrenia-like sensory phenotypes in Setd1a+/- mice. Biol. Psychiatry 88, 215–223 (2020).

62. D. P. Leone, K. Srinivasan, B. Chen, E. Alcamo, S. K. McConnell, The determination of projection neuron identity in the developing cerebral cortex. Curr. Opin. Neurobiol. 18, 28–35 (2008).

63. L. Roseman, L. Demetriou, M. B. Wall, D. J. Nutt, R. L. Carhart-Harris, Increased amygdala responses to emotional faces after psilocybin for treatment-resistant depression. Neuropharmacology 142, 263–269 (2018).

64. M. K. Doss, M. Považan, M. D. Rosenberg, N. D. Sepeda, A. K. Davis, P. H. Finan, G. S. Smith, J. J. Pekar, P. B. Barker, R. R. Griffiths, F. S. Barrett, Psilocybin therapy increases cognitive and neural flexibility in patients with major depressive disorder. Transl. Psychiatry 11, 574 (2021).

65. T. Kube, R. Schwarting, L. Rozenkrantz, J. A. Glombiewski, W. Rief, Distorted cognitive processes in major depression: A predictive processing perspective. Biol. Psychiatry 87, 388–398 (2020).

66. H.-W. Shen, X.-L. Jiang, J. C. Winter, A.-M. Yu, Psychedelic 5-methoxy-N,N-dimethyltryptamine: metabolism, pharmacokinetics, drug interactions, and pharmacological actions. Curr. Drug Metab. 11, 659–666 (2010).

67. S. J. Jefferson, I. Gregg, M. Dibbs, C. Liao, H. Wu, P. A. Davoudian, S. C. Woodburn, P. H. Wehrle, J. S. Sprouse, A. M. Sherwood, A. P. Kaye, C. Pittenger, A. C. Kwan, 5-MeO-DMT modifies innate behaviors and promotes structural neural plasticity in mice. Neuropsychopharmacology 48, 1257–1266 (2023).

68. A. Delorme, S. Makeig, EEGLAB: an open source toolbox for analysis of single-trial EEG dynamics including independent component analysis. J. Neurosci. Methods 134, 9–21 (2004).

69. S. Moratti, B. A. Clementz, Y. Gao, T. Ortiz, A. Keil, Neural mechanisms of evoked oscillations: stability and interaction with transient events. Hum. Brain Mapp. 28, 1318–1333 (2007).

70. M. C. Dadarlat, M. P. Stryker, Locomotion enhances neural encoding of visual stimuli in mouse V1. J. Neurosci. 37, 3764–3775 (2017).

71. C. G. Gallimore, D. A. Ricci, J. P. Hamm, Spatiotemporal dynamics across visual cortical laminae support a predictive coding framework for interpreting mismatch responses. Cereb. Cortex 33, 9417–9428 (2023).

72. A. M. Bastos, M. Lundqvist, A. S. Waite, N. Kopell, E. K. Miller, Layer and rhythm specificity for predictive routing. Proc. Natl. Acad. Sci. U. S. A. 117, 31459–31469 (2020).

73. R. Oostenveld, P. Fries, E. Maris, J.-M. Schoffelen, FieldTrip: Open source software for advanced analysis of MEG, EEG, and invasive electrophysiological data. Comput. Intell. Neurosci. 2011, 156869 (2011).

74. J. Jackson, M. M. Karnani, B. V. Zemelman, D. Burdakov, A. K. Lee, Inhibitory control of prefrontal cortex by the claustrum. Neuron 99, 1029–1039.e4 (2018).

75. Y. K. Hong, C. O. Lacefield, C. C. Rodgers, R. M. Bruno, Sensation, movement and learning in the absence of barrel cortex. Nature 561, 542–546 (2018).

76. G. T. Prusky, R. M. Douglas, Characterization of mouse cortical spatial vision. Vision Res. 44, 3411–3418 (2004).

