## Supplemental methods and figures for "Psychedelics relax predictive processing in the post-acute period by remodeling cortico-cortical feedback circuits"

**Supplemental materials.**

**Supplemental Methods:**

This section provides an overview of the supplemental behavioral assessments used to compare our experimental groups. We focused on personality factors and risk-taking behaviors. Each participant was administered a computerized version of the Domain-Specific Risk-Taking (DOSPERT) scale and the OCEAN personality assessment following the EEG data collection. All subjects were administered both the DOSPERT and the OCEAN, except for three in the 5-MeO-DMT group. Statistics in the DOSPERT/OCEAN portion of table S1 reflect n=12 for that group.

**DOSPERT Scale**

The 30-item DOSPERT scale, a comprehensive tool designed to evaluate risk-taking attitudes and behaviors across different domains (e.g., financial, health/safety, recreational, ethical, and social), was utilized to assess participants' propensity for risk-taking. This scale comprises 6 items, rated on a Likert scale, for each domain. For each item, participants were asked the likelihood that they would engage in the activity on a seven-point scale, ranging from extremely unlikely (with a numeric value of 1) to extremely likely (with a numeric value of 7). Responses were summed across each domain to calculate domain scores for risk-taking behavior

Two-sample independent t-tests were used to investigate potential differences in risk-taking behavior between our control group (CNT) and recent 5-HT2A psychedelic users (PSY). We identified only a difference in Health/Safety risk taking between CNT and PSY (Table 1). We also repeated this for the 5-MeO-DMT group, comparing to CNT. No group differences were identified (Table S1).

**OCEAN Model Assessment**

A 50-item survey version of the five factor model of personality(*33*), also known as the OCEAN model (Openness, Conscientiousness, Extraversion, Agreeableness, and Neuroticism) was employed to assess for group differences in personality traits of our participants. This model is well established and provides a robust framework for understanding personality.

Two-sample independent t-tests were used to investigate potential differences in each of the 5 factors between our control group (CNT) and recent 5-HT2A psychedelic users (PSY). We identified no significant group differences between CNT and PSY (Table 1). We also repeated this for the 5-MeO-DMT group, comparing to CNT. We found a modest group difference in openness, with the 5-MeO-DMT group showing slightly more openness to experience (Table S1).

**Paradigm choice in mice**

Humans completed a saccadic prediction task in order to glean neural and behavioral indices of predictive processing in a quickly learned, rapidly acquired paradigm (<10 minutes) that manipulates predictability (target appearing at standard or new location) while holding salience constant (i.e., a suddenly occurring stimulus after 1 second of blank screen and requiring an eye movement). Because predictive suppression was found to differ between groups, we followed up our human findings in mice using a combination of global-local oddball and standard oddball paradigms, as it involves rapidly presented stimuli that, like the human study, are also equally salient (i.e., the “B” stimulus, which is 90-degrees of orientation different from the stimulus encountered 500ms prior) but *differentially predictable* in both settings. This shift was thus able to focus on a similar function (predictive suppression and deviance detection), but addressed two issues with cross species translation. First, our mouse experiments were passive. Training mice to associate a motor output with a sensory input requires days to weeks of training, and the role of sensory cortex in such learned behaviors is ultimately uncertain (*75*). Second, as mice do not foveate and generally have less acuity (but more sensitivity to low-spatial frequencies and motion (*76*)), the same “star” stimuli used in the human study would have been inappropriate. Generally, given the differences in the visual and motor systems (i.e., mice do not actively saccade) between mice and humans, equivalence of stimulation is practically unattainable, even if all physical stimulus features are held constant. Nonetheless, we did record EEG responses to a basic oddball paradigm in humans (Supplemental Fig. S8) and, like psilocybin treated mice, human deviance detection (greater responses to deviant vs control) was intact after psychedelic use

|  | **Control Group** | **5-MeO-DMT Users** | | **Statistics** | **p-value** | |
| --- | --- | --- | --- | --- | --- | --- |
| Subjects | 16 | 15 | | - | | - |
| Age (years) | 37.44 ± 15.082 | 45.67 ± 11.1 | | t(26) = 1.71 | | *p* = 0.10 |
| Sex | 6 Male; 10 Female | 6 Male; 9 Female | | *c*^2^(1)= 0.020 | | *p* = 0.866 |
| Race | 11 White (non-Hispanic)  2 Hispanic or Latino  2 Black or African American  1 Asian | 10 White (non-Hispanic)  4 Hispanic or Latino  1 Black or African American  0 Asian | | *c*^2^(3)= 2.017 | | *p* = 0.569 |
| Education Level | 3 High School Degree  8 Associate or Bachelor's Degree  5 Advanced Degree | 3 High School Degree  6 Associate or Bachelor's Degree  6 Advanced Degree | | *c*^2^(5)=6.441 | | *p* = 0.266 |
| Marijuana Use | 15 Never Used / Infrequent User  1 Frequent User | 11 Never Used / Infrequent User  4 Frequent User | | *c*^2^(3)= 3.104 | | *p* = 0.376 |
| Days Since Last Psychedelic Use | - | 15.2 ± 8.01 (range: 2-27) | | - | | - |
| **OCEAN Assessment Personality Traits** | | | | | | |
| Openness | 29.56 ± 5.240 | 33.75 ± 3.571 | | t(26) = 2.379 | | *p*= 0.025* |
| Contentiousness | 27.87 ± 8.461 | 27.17 ± 10.143 | | t(26) = -0.201 | | *p* = 0.842 |
| Extraversion | 23.06 ± 7.550 | 26.58 ± 6.708 | | t(26) = 1.280 | | *p* = 0.212 |
| Agreeableness | 34.12 ± 3.500 | 32.42 ± 4.122 | | t(26) = -1.185 | | *p* = 0.247 |
| Neuroticism | 24.68 ± 7.631 | 25.91 ± 8.928 | | t(26) = 0.528 | | *p* = 0.698 |
| **DOSPERT Risk-Taking Domains** | | | | | | |
| Ethical | 11.25 ± 3.715 | 10.33 ± 4.097 | | t(26) = 0.618 | | *p* = 0.542 |
| Financial | 15.12 ± 7.032 | 13.68 ± 5.331 | | t(26) = 0.600 | | *p* = 0.554 |
| Health/Safety | 18.25 ± 6.266 | 15.75± 7.200 | | t(26) = -0.980 | | *p* = 0.336 |
| Recreational | 22.56 ± 9.654 | 23.92± 10.004 | | t(26) = 0.362 | | *p* = 0.720 |
| Social | 33.56 ± 4.953 | 34.58± 5.089 | | t(26) = 0.533 | | *p* = 0.598 |

**Table S1: Demographic data, personality scores, and risk-taking among control and recent psychedelics-users.** Group values represent means, +/- standard deviations.

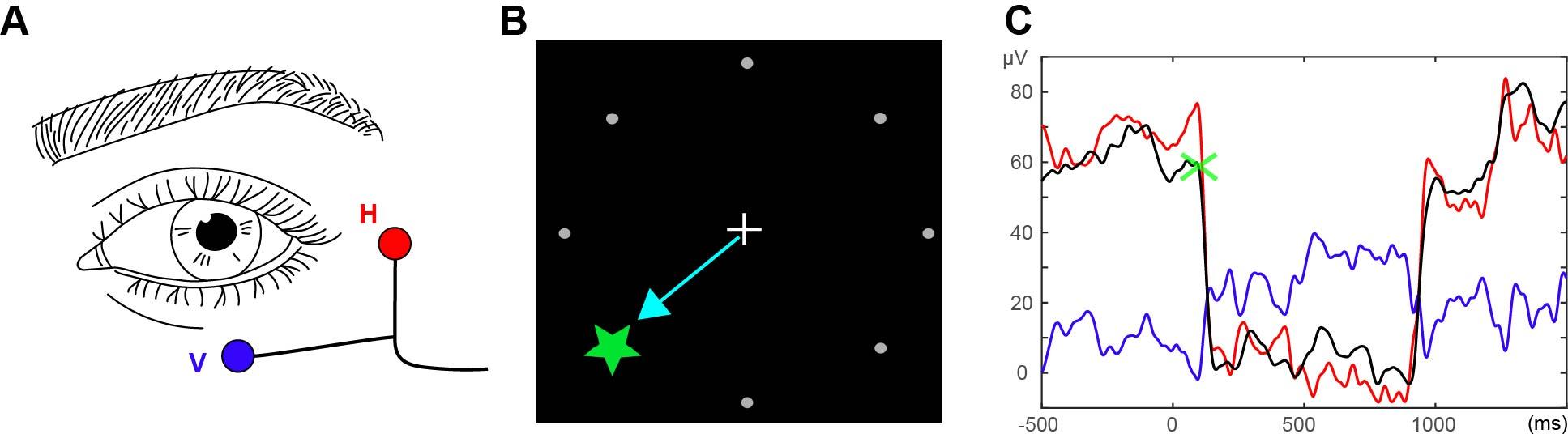

**Supplemental figure 1: Electrooculography (EOG).** A) Surface electrodes were placed <1cm to the left of the outer canthi of the left eye (horizontal: H) and <1cm below the left eye (vertical: V). B) example trial, where a stimulus appears in one of 8 locations. C) Example EOG from one subject. The sum of the absolute values (black) of the two EOGs (red: H, blue: V) were plotted for each trial and used to score the onset of each saccade (green X).

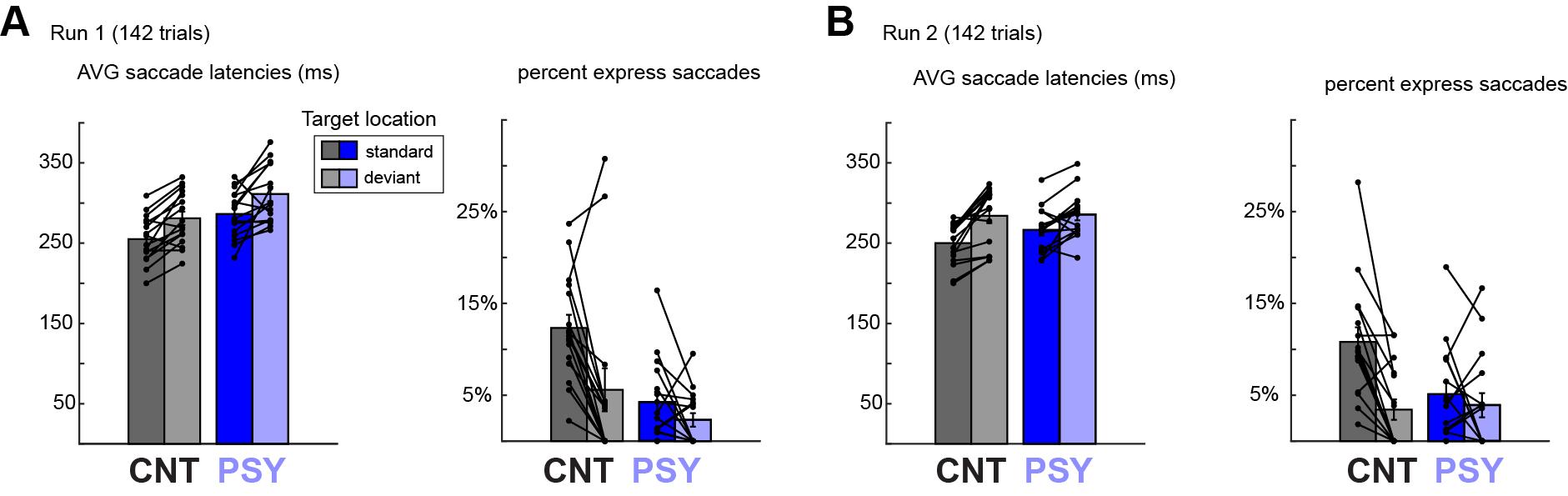

**Supplemental figure 2: Predictive saccadic effects were stable between two subsequent runs of the SPT**. A,B) all subjects completed two runs of the saccadic prediction task. Similar effects of overall average latency were seen in both runs. However, a trend toward a group by stimulus location was seen in run two (F(1,30)=3.38, p=.076, with PSY showing slightly less of a difference between standard (predictable) and deviant trials. For proportion of express saccades, a group by stimulus location interaction was seen in both runs (run1-F^interaction^(1,30)=4.87, p=0.035; run2-F^interaction^(1,30)=9.90, p=0.004).

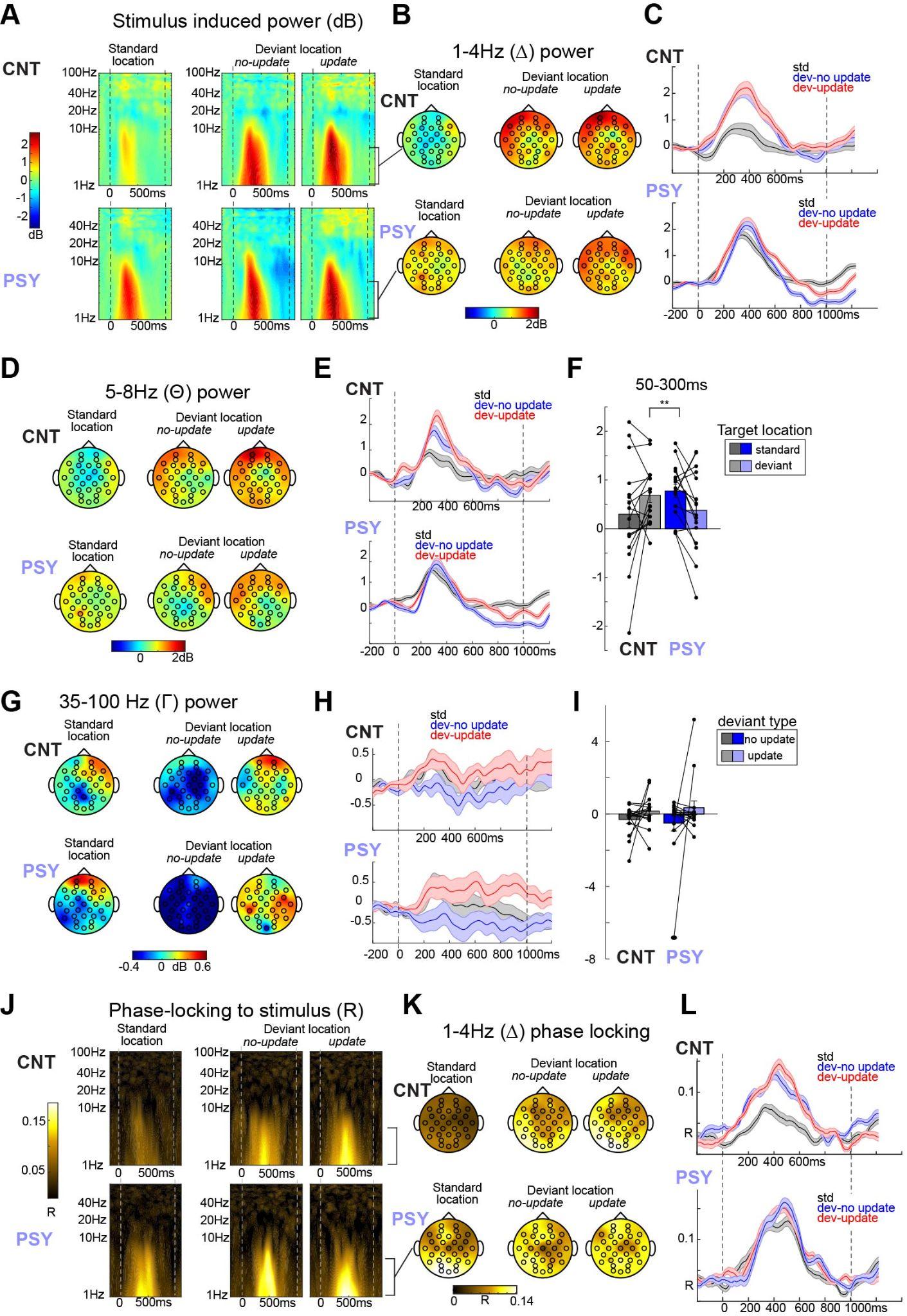

**Supplemental figure 3: Effects of updating and other frequency band effects during the Saccadic Prediction Task.** For power, no main effects of deviant type or deviant type by group interactions were present for any frequency band or time-bin (early, late), although a trend-level effect on gamma power for deviant type was present at p=.11 (G-I). A) Stimulus induced power spectra, separated by deviant type (update vs no-update). B) Delta power (50-600ms) plotted across scalp, separated by deviant type. C) Delta power across all electrodes, separated by deviant type. D) Theta power plotted across the scalp, and E) averaged across electrodes and plotted as a function of time. No effect of deviant type was present, but a F) group by stimulus location (deviant vs predictable) was present for early latencies (50-300ms), matching the delta effects seen in figure 2. G-H) same as D-E, but for Gamma power. (I) is similar to (F) but compares between deviant types. J-L) Same as A-C, but for phase-locking. For phase-locking, no main effects of deviant type or deviant type by group interactions were present for any frequency band or time-bin (early, late).

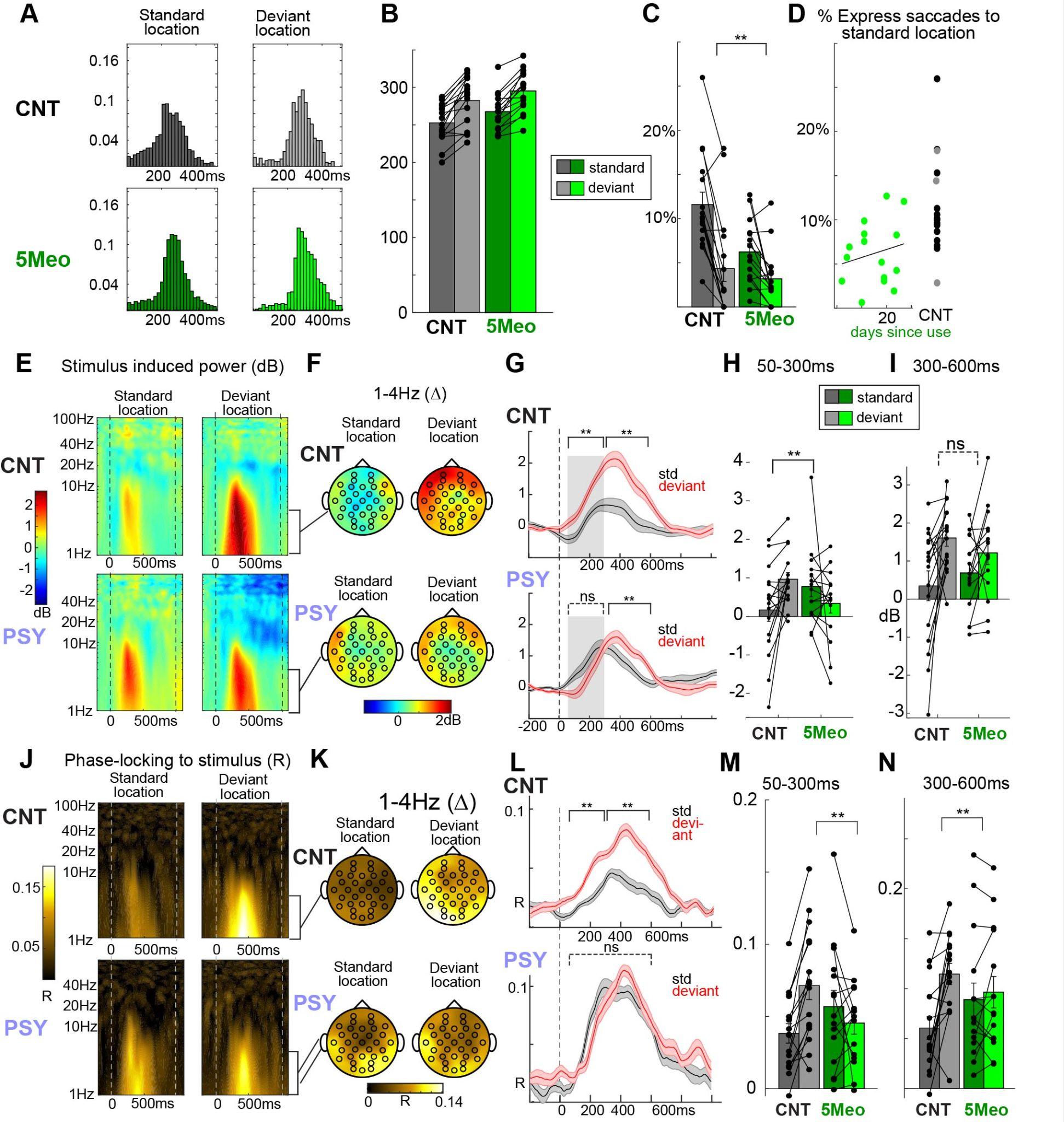

**Supplemental figure 4: cohort 2 using 5-Meo-DMT.** A) A) Fifteen participants who recently took 5-MeO-DMT completed the SPT task and displayed saccadic latency distributions distinct from control subjects (CNT) to standard (predictable) targets, but, to deviant targets, more similar to CNT. B) Mean latency and C) percent express saccade bar plots, each dot is one subject’s average. The 5-MeO-DMT displayed fewer express saccades to predictable locations than CNT. D) Proportion of express saccades showed a trend-level correlation with number of days since taking the dose (r=.33, p=.11 one-tailed). E) Stimulus induced power spectra and F) scalp topographies of 50 to 600ms delta (1-4Hz) power for each group. G) Stimulus induced power to the onset of targets in the predictable location was not smaller than to targets in the deviant locations in the 5-MeO group for H) early (pre/perisaccade activity; delta-band), but I) it was for the later time ranges (mirroring effects seen in PSY (figure 2)). J-N) same as G-H, but for phase locking factor to stimulus onset. **p<.01, *p<.05.

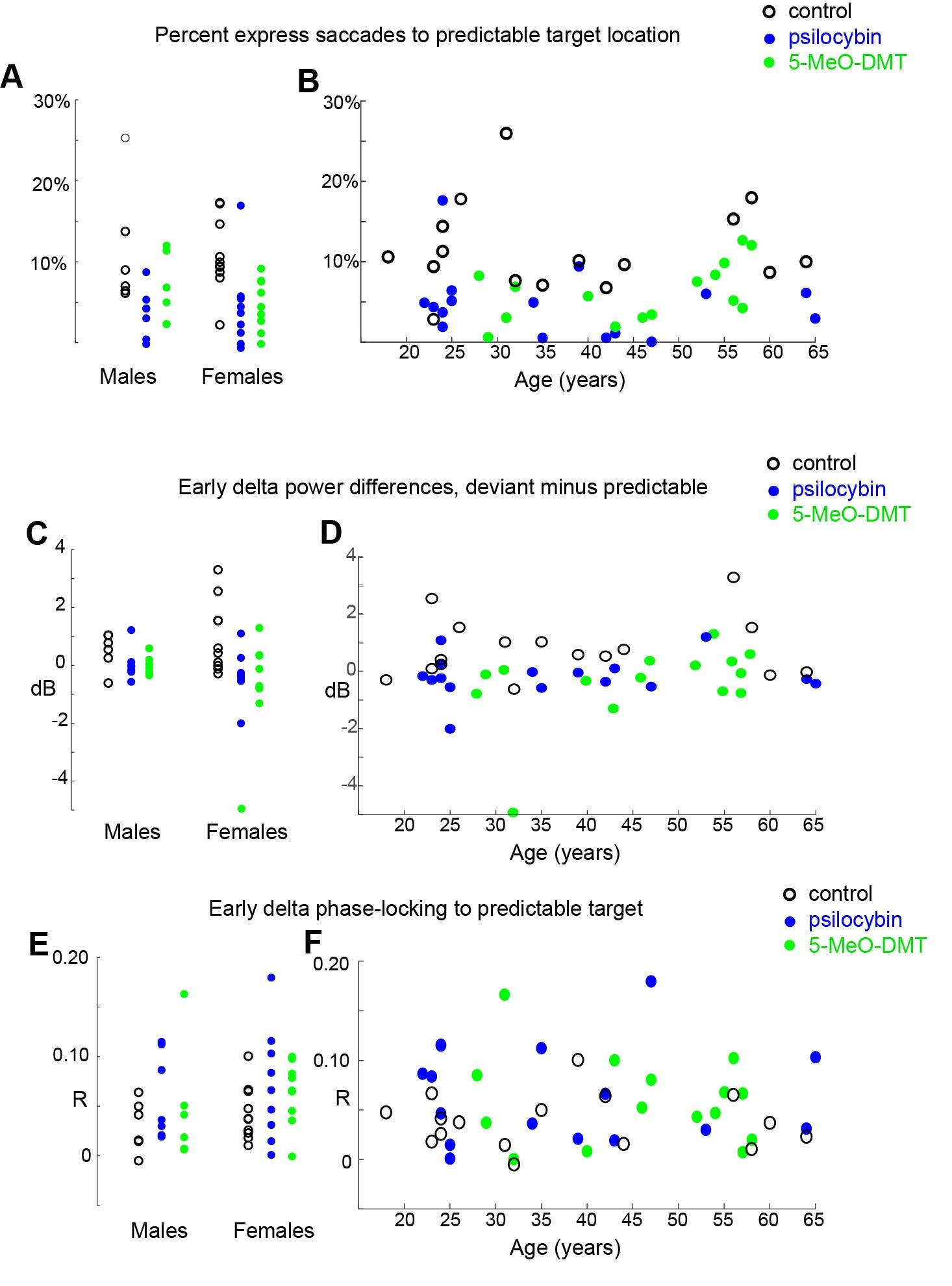

**Supplemental figure 5: Age and sex effects.** A) Percent of saccades that were express to the predictable target plotted as a function of sex and B) age for each group. C,D) same as A,B for delta power (all electrodes) to the onset (50-300ms) of predictable vs deviant targets. E-F) same as E,F but for phase locking factor to the onset of the predictable target. We chose to focus on differences for C,D because power varies more widely across subjects than relatively absolute measures like saccade percentages or phase locking (which are both bound between 0 and 1).

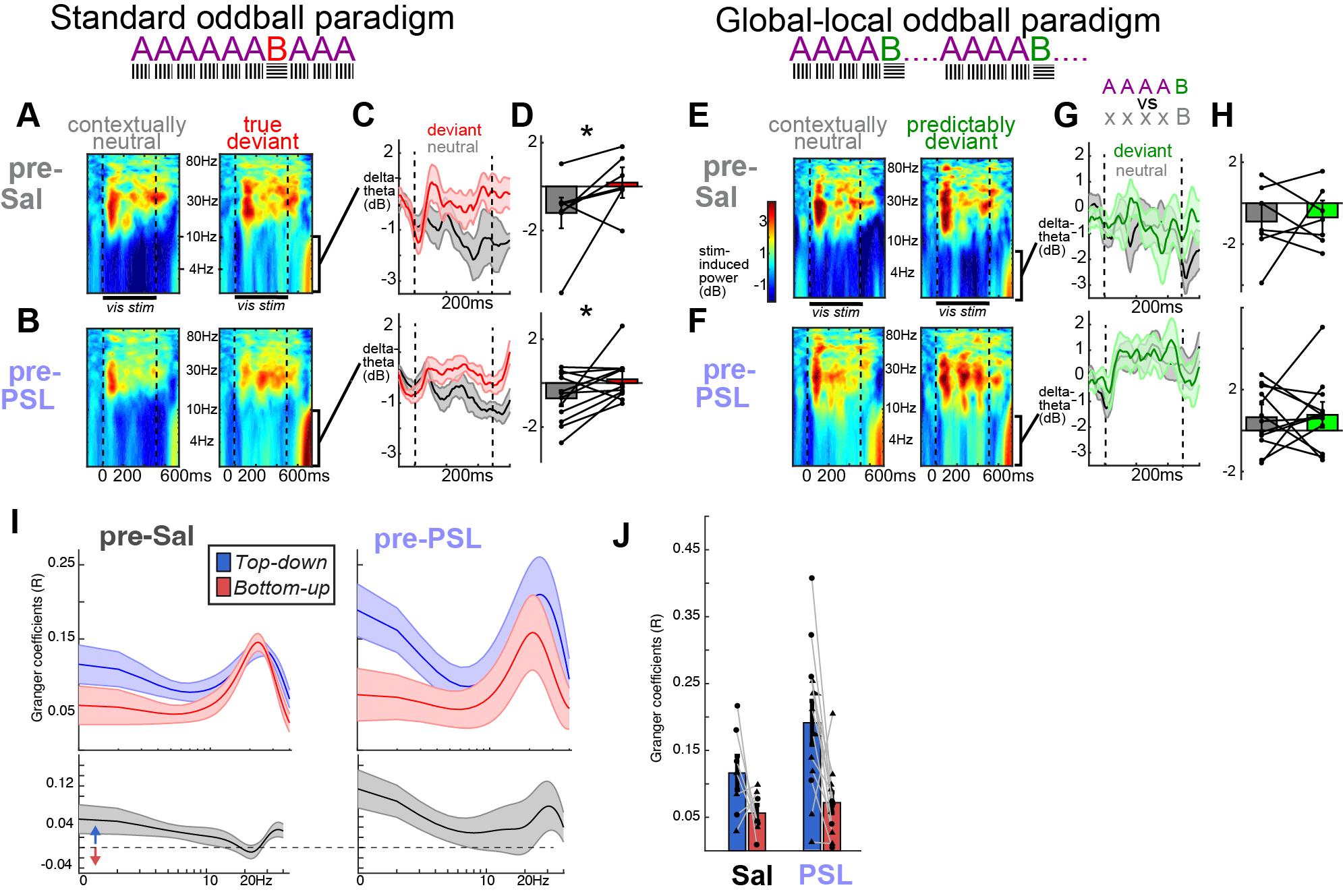

**Supplemental figure 6: Baseline analysis for global-local.** Matching figure 3, but for pre-treatment baseline. One hour prior to treatment with saline or psilocybin, mice were exposed to the standard oddball paradigm and the global local oddball paradigm. No group by context (deviant vs control) by paradigm interaction was present F(1,67)=0.149, p=.699. A) Time-frequency stimulus-induced power plots demonstrate significant deviance detection to “B” stimulus in the standard oddball paradigm in both saline mice or B) mice to be treated later with psilocybin. C) Averaged power from 2-10Hz and D) again averaged from 50-600ms post stimulus onset (F^context^(1,33)=6.65, p=.014; t(7)=1.97, p=.048 saline; t(11)=2.39, p=.018 psilocybin). E-H) Same as A-D, but for the global-local oddball paradigm, in which the “B” stimulus is predictable. No deviance detection was observed in this period for either group (F^context^(1,33)=0.0688, p=.795). *p<.05 paired t-tests, one tailed. I) Top- average spectra of top-down and bottom-up Granger coefficients (directed influence from ACa to V1 or V1 to ACa, respectively) for saline control (left) vs psychedelic groups (right). Bottom - top-down minus bottom-up spectra. j) Average GC values for all stimulus/ITI time-bins, for 2-4Hz. Each dot is one mouse. Triangles are males. No significant interactions involving group and direction were observed at this baseline measurement. Since you want PSY for in humans and PSL in mice- change in figure

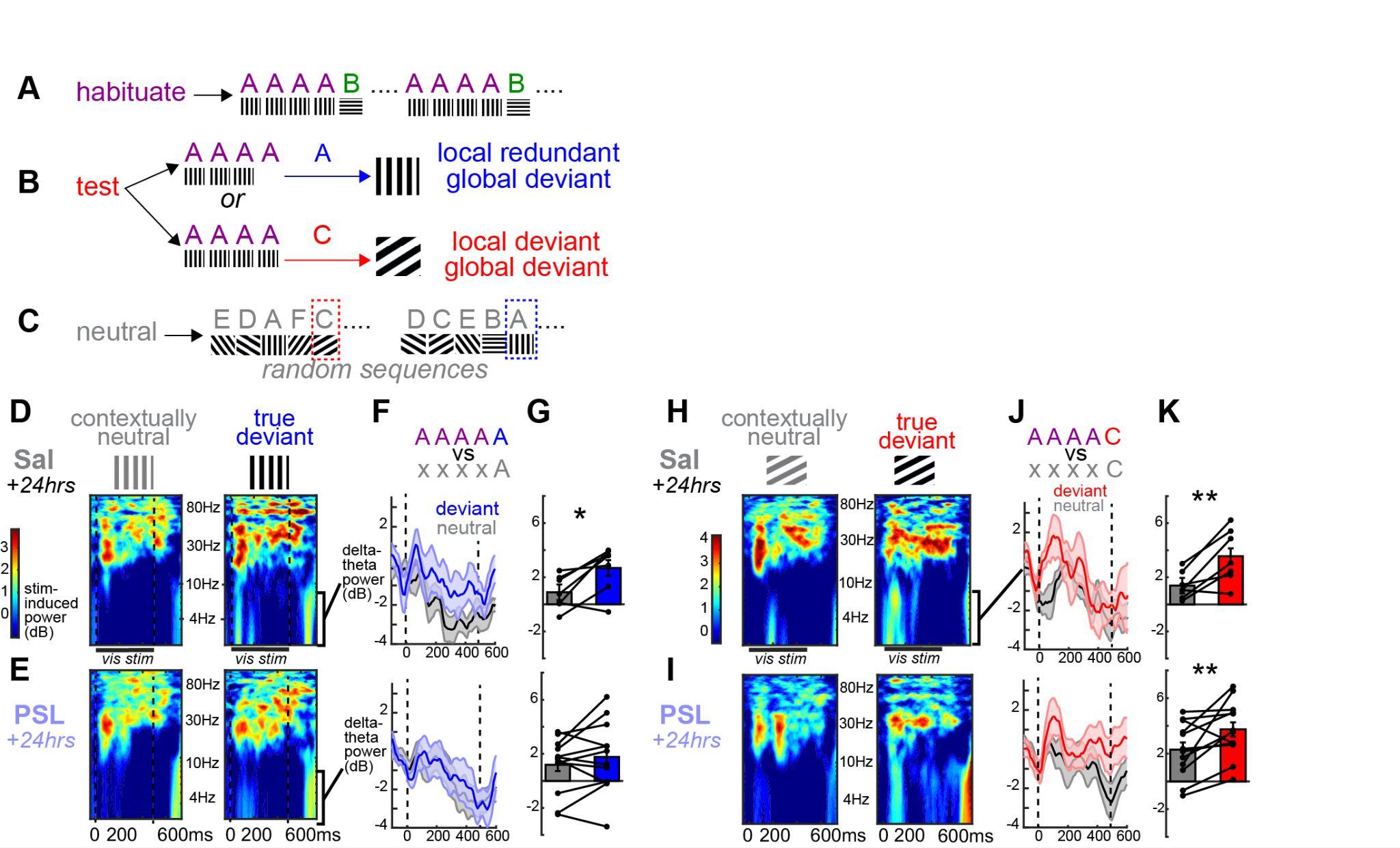

**Supplemental figure 7. Catch trials during global-local oddball at 24-hrs post dose.** A) During the global-local oddball paradigm, mice are habituated to 5 stimulus sequences of an A orientation 4 times (90 deg) followed by a B orientation (0 deg; deviant), with 500ms stimulus duration, 500ms inter-stimulus interval, 5-6 second inter-trial interval. B) During the test run, one in every 6 trials ended in a deviant – either a fifth A stimulus (which was locally redundant with the previous stimuli, but globally deviant) or a C stimulus (halfway between A and B – 45 degrees; locally deviant and globally deviant). C) Responses were contrasted to a neutral control, where one of 8 random orientation was presented. D) Power spectra for the saline treated and E) mice 24 hours after treatment for the 90 degree stimulus during the control run and when it was the local redundant but global deviant (AAAA-A). F) Line plots of delta-theta power to each condition and G) max delta-theta power during the stimulus interval (0 to 500ms). Significant DD in the saline but not the psilocybin group. H-K) same as D-G, but for the AAAAC (local and global deviant). Analysis note -- deviance detection on these trials (control vs oddball) was somewhat weaker than observed during the standard oddball for both groups, perhaps due to lower signal to noise ratio (only 6 trials) or because the difference between the current and immediate previous stimulus was smaller (45 or 0 deg difference, vs 90 deg difference in the standard oddball). So, for both groups, we analysed the maximum response post-stimulus (0-500ms post stimulus onset), rather than the mean. This did not change the pattern of effects, but helped to reveal statistical significance. *p<.05; **p<.01. one tailed paired samples t-tests.

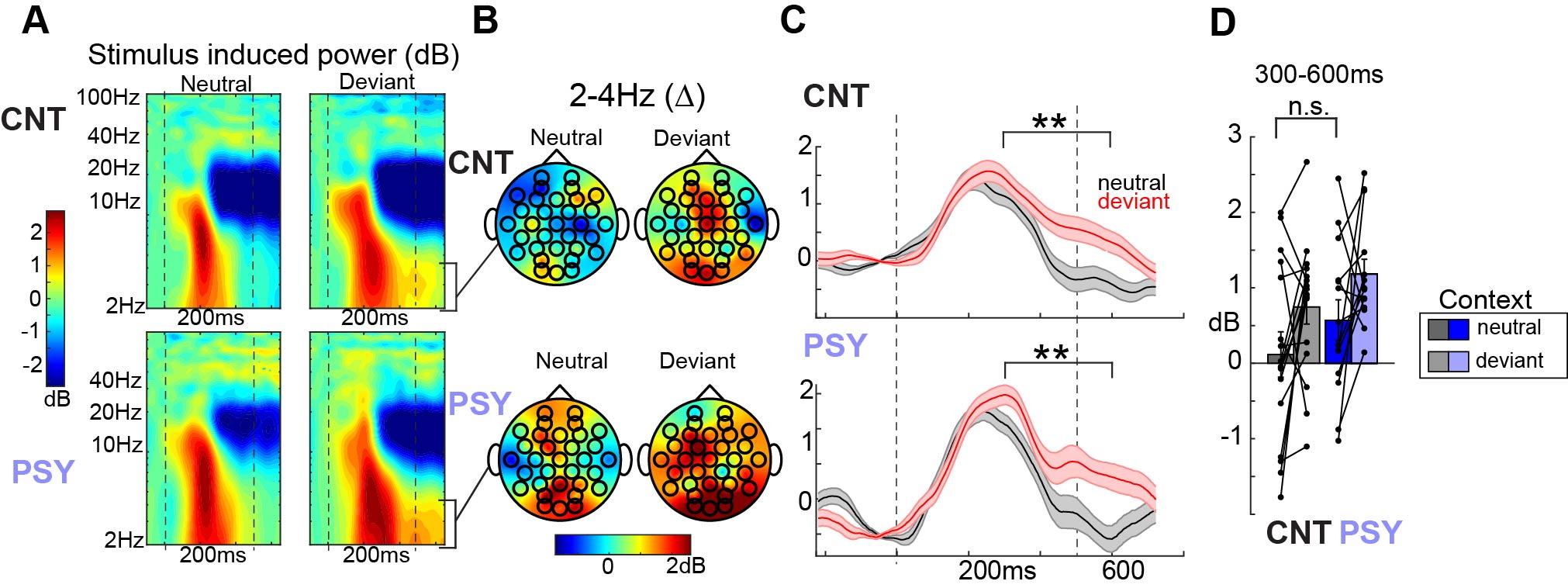

**Supplemental figure 8. Standard oddball in human participants.** A) Time-frequency plots of stimulus-induced delta power, averaged across all electrodes, to oriented bar stimuli presented during a many-standards control sequence (stimulus is neutral in context) vs during an oddball sequence (stimulus is deviant in context). B) Topographies for each group and condition of delta power averaged from 50 to 600ms post stimulus onset. C) Averaged delta power (2-4Hz) averaged across occipital-parietal electrodes plotted as a function of time for control subjects and subjects taking psychedelics. Only the late period (300 – 600ms post stimulus onset) showed significant deviance detection, regardless of group. D) Values from B, averaged from 300 to 600ms post stimulus onset. Across all subjects, an effect of context was observed (deviant vs control; F(1,28)=6.92, p=.0137). No group by context interaction was present (F(1,28)=0.00, p=.976) as in the mouse experiments. For this portion, the same parameters and analyses were used as in the mouse experiments, except we focused only on the delta-band (1-4Hz) and later time-bin (300-600ms), as no deviance detection was observed early or in theta. The visual oddball paradigm was presented using Psychopy in a darkened room where participants were seated 100cm away from an LCD monitor (19-27 inches, 60Hz refresh rate). Square-wave gratings (12.75”x12.75” squares; 18.18 degrees of visual angle) were presented at 100% contrast and 2.0 cycles per degree, drifting at two cycles per second lasting 500ms, with a randomly jittered 450-550ms inter stimulus interval (black screen). We presented three separate sequences – each consisting of 200 trials and lasting ≈200 seconds.
